# Dendrites use mechanosensitive channels to proofread ligand-mediated guidance during morphogenesis

**DOI:** 10.1101/2022.02.28.482179

**Authors:** Li Tao, Sean Coakley, Rebecca Shi, Kang Shen

## Abstract

Ligand-receptor interactions guide axon navigation and dendrite arborization. Mechanical forces also influence guidance choices. However, the nature of such mechanical stimulations, the mechanosensors identity and how they interact with guidance receptors are unknown. Here, we demonstrate that mechanosensitive DEG/ENaC channels are required for dendritic arbor morphogenesis in *Caenorhabditis elegans*. Inhibition of DEG/ENaC channels causes reduced dendritic outgrowth and branching *in vivo*, a phenotype that is alleviated by overexpression of the mechanosensitive channels PEZO-1/Piezo or YVC1/TrpY1. DEG/ENaCs trigger local Ca^2+^ transients in growing dendritic filopodia via activation of L-type voltage-gated Ca^2+^ channels. Anchoring of filopodia by guidance ligand/receptor is required for the mechanical activation of DEG/ENaC channels. Therefore, mechanosensitive channels serve as a checkpoint for appropriate chemoaffinity by activating Ca^2+^ transients required for neurite growth.

## Introduction

Mechanosensitive ion channels play essential roles in the sensory transduction of mechanical force and underpin sensations of touch and hearing. Several classes of mechanosensitive channels have been identified and described to transduce physical stimulus into changes in membrane potential: the degenerin and epithelial Na^+^ channels (DEG/ENaC), transient receptor potential (TRP) proteins, transmembrane channel-like (TMC) proteins, mechanosensitive K^+^ channels and Piezo ion channels (Árnadóttir and Chalfie, 2010; Geffeney and Goodman, 2012; Ranade et al., 2015; Venkatachalam and Montell, 2007). Unlike receptors in other sensory systems, mechanosensors are broadly expressed outside of sensory systems and play unexpected roles in development (Ranade et al., 2015). For example, the mechanically-gated ion channel Piezo1 is activated in endothelial cells by blood flow-mediated sheer force, and is required to trigger Ca^2+^ influx in endothelial cells for subsequent cellular differentiation (Li et al., 2014). The surprising ability of mechanosensitive channels to respond to mechanical stimuli in various tissues throughout the body supports the notion that mechanotransduction pathways may regulate numerous developmental processes.

The involvement of mechanosensitive channels in neuronal development is not understood. The wiring of the nervous system is a complex process instructed by genetic cues and influenced by environmental signals. Both evoked and spontaneous neuronal activity play essential roles in the wiring of neural circuits (Pan and Monje, 2020). At a cellular level, developing growth cones are highly sensitive to changes Ca^2+^ concentration (Gasperini et al., 2017) and the spatial localization and amplitude of Ca^2+^ can influence growth cone extension and direct neurite outgrowth (Kater and Mills, 1991; Tojima et al., 2011; Wen et al., 2004; Zheng, 2000). At a molecular level, diffusible and membrane tethered ligands and their corresponding membrane receptors guide axon navigation and dendrite branching (Blockus and Chédotal, 2016; Onishi et al., 2014; Pasterkamp, 2012). Importantly, several guidance cues, including netrin-1 (Hong et al., 2000), neurotrophins (Wang and Zheng, 1998), and semaphorins (Behar et al., 1999), can modulate Ca^2+^ dynamics in growth cones, suggesting that molecular guidance pathways intersect with activity dependent mechanisms.

While a large body of work demonstrates that growth cones sense chemical signals, emerging evidence suggests that they might also respond to physical forces (Kerstein et al., 2015). The rigidity of substrates can also influence axonal growth cone behavior (Flanagan et al., 2002; Koch et al., 2012; Koser et al., 2016). Recent evidence shows that chemical ligand can also provide as mechanical cues. For example, physical attachment of Netrin to its substrate plays important roles in netrin’s effects on growth cones *in vitro*. More directly, the authors showed that restraining netrin coated beads to reach a certain tension between the beads and growth cones increased netrin signaling (Moore et al., 2012). During axonal extension, traction forces are generated when membrane receptors are coupled to the non-muscle myosin II-mediated retrograde actin flow (Brown and Bridgman, 2003; Suter and Forscher, 2000). Together, these data suggest that mechanical force related to neurite outgrowth could be a source of neuronal activity with the ability to modulate axonal guidance. However, the *in vivo* relevance of this model and the potential involvement of mechanosensitive ion channels remains elusive.

In this study, we demonstrate that proper guidance ligand-receptor interactions activate mechanosensitive ion channels to regulate the growth and branching of dendrites *in vivo*. We observed frequent local Ca^2+^ transients in outgrowing dendritic filopodia, which are dependent on the DEG/ENaC family of mechanosensitive ion channels and L-type voltage-gated Ca^2+^ channels (VGCCs), indicating that the forces incurred at the growing dendrite can activate mechanosensitive ion channels. Activation of DEG/ENaCs requires dendrite guidance ligands and the proper actin cytoskeleton organization. Together, our findings suggest that this mechanosensitive property of the growth cone acts as a proof-reading mechanism for the correct guidance and outgrowth of dendrites.

## Results

### Local Ca^2+^ transients play important roles in dendrite formation

To understand neuronal activity during morphogenesis, we monitored spontaneous Ca^2+^ activity of the PVD neuron during the outgrowth of terminal 4° dendritic branches and observed distinct types of Ca^2+^ transients. These included frequent local Ca^2+^ transients restricted to individual 4° and 3° dendritic branches and occasional global Ca^2+^ transients, which propagate throughout one or several menorah structures (Fig. 1, A and B). Within the local Ca^2+^ transients, the vast majority were restricted to the 4° terminal dendritic branches during their outgrowth (Fig. 1C and Movie S1). These local 4° Ca^2+^ transients lasted for 5-15s and many initiated from the tips of 4° dendrites (Fig. 1B and fig. S1). 39% of the 4° Ca^2+^ transients are restricted to the tips throughout the frames, while 26% propagate to the entire 4° branches. The remaining 35% further propagate to 3°, but not 2° branches, forming an L-shape signal (fig. S1). A small number of Ca^2+^ transients occurred in 3° or the entire menorah (Fig. 1, B and C). We have previously shown that those global, menorah wide Ca^2+^ transients are triggered by body movement (Tao et al., 2019).

**Figure 1.**
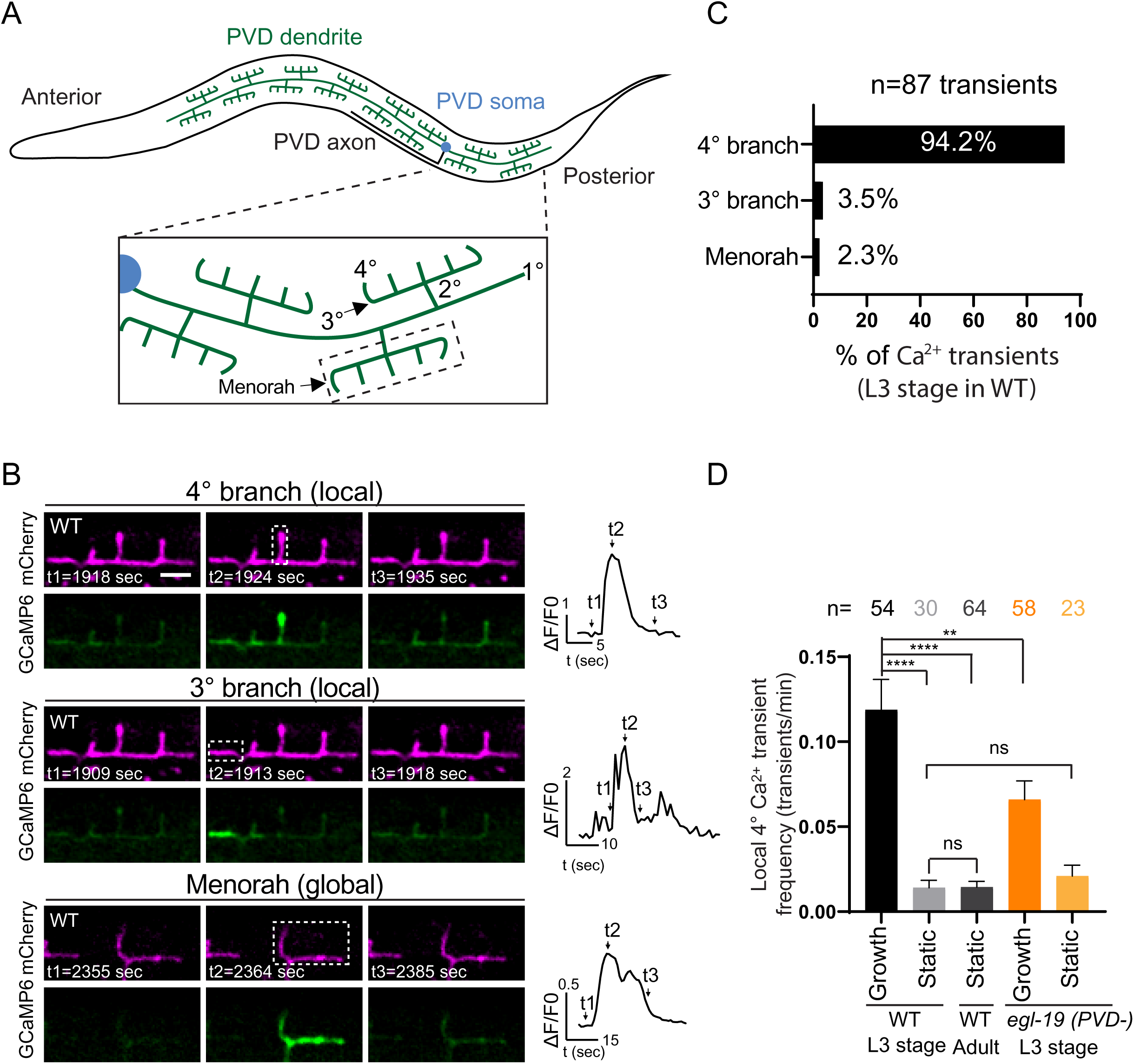
Local calcium transients occur during 4° dendrite growth. (**A**) Scheme showing the morphology of the PVD neuron in *C. elegans*. (**B**) Confocal images of GCaMP6 (green), mCherry (magenta) and corresponding Ca^2+^ traces, showing different types of calcium transients observed during development in wild-type (WT) late L3 animals. Scale bar: 5 μm. Rectangles: Regions for GCaMP6 intensity measurement. (**C**) Percentage of each type of calcium transient observed during 4° dendrite growth. 4 independent animals were imaged. (**D**) Quantification of the local Ca^2+^ transient frequency (number of events per minute) in growing and static 4° dendrites in WT and *egl-19* PVD knockdown *(PVD-)* animals in late L3 stage animals and in mature 4° dendrites (static) in young adult animals. Error bars: SEM. Statistics, One-way ANOVA with a Tukey correction. ****p<0.0001. **p<0.01. ns: not significant. 6 independent animals were used for each genotype.

Importantly, local 4° Ca^2+^ transients were much more frequent during the outgrowth of 4° branches (“growth” in Fig. 1D), compared to branches that were not growing (“static” in Fig. 1D), indicating that these local Ca^2+^ events might be related to dendrite outgrowth. Consistently, these local 4° Ca^2+^ transients were rare in adult animals where dendrites stopped growing (Fig. 1D). PVD cell-specific knockdown of the α-subunit of the L-type VGCC, *egl-19*, dramatically reduced the local 4° Ca^2+^ transient frequency in the growing 4° branches, indicating that activation of L-type VGCCs during dendrite terminal outgrowth gives rise to local Ca^2+^ transients (Fig. 1D).

To ask whether the Ca^2+^ transients are necessary for dendrite outgrowth, we examined the PVD dendrite arbor in the *egl-19* PVD cell-specific knockdown animals. The knockdown animals showed a loss of 4° branches (Fig. 2, A and B). To further characterize dendrite formation, we performed time-lapse recording of outgrowth events. In wild-type animals, the PVD 4° branches initiate as filopodia and exhibit continuous growth until reaching the dorsal or ventral nerve cord. The *egl-19* knockdown PVDs could initiate dendritic filopodia, but failed to extend these filopodia beyond ∼1 µm, leading to slowed overall growth speed and shortened dendrites (Fig. 2, C-F and Movie S2-S3). Null mutants of the N-type VGCC, *unc-2*, showed normal dendrite morphogenesis, indicating that the L-type VGCCs are specifically involved in dendrite growth (Fig. 2, A and B).

**Figure 2.**
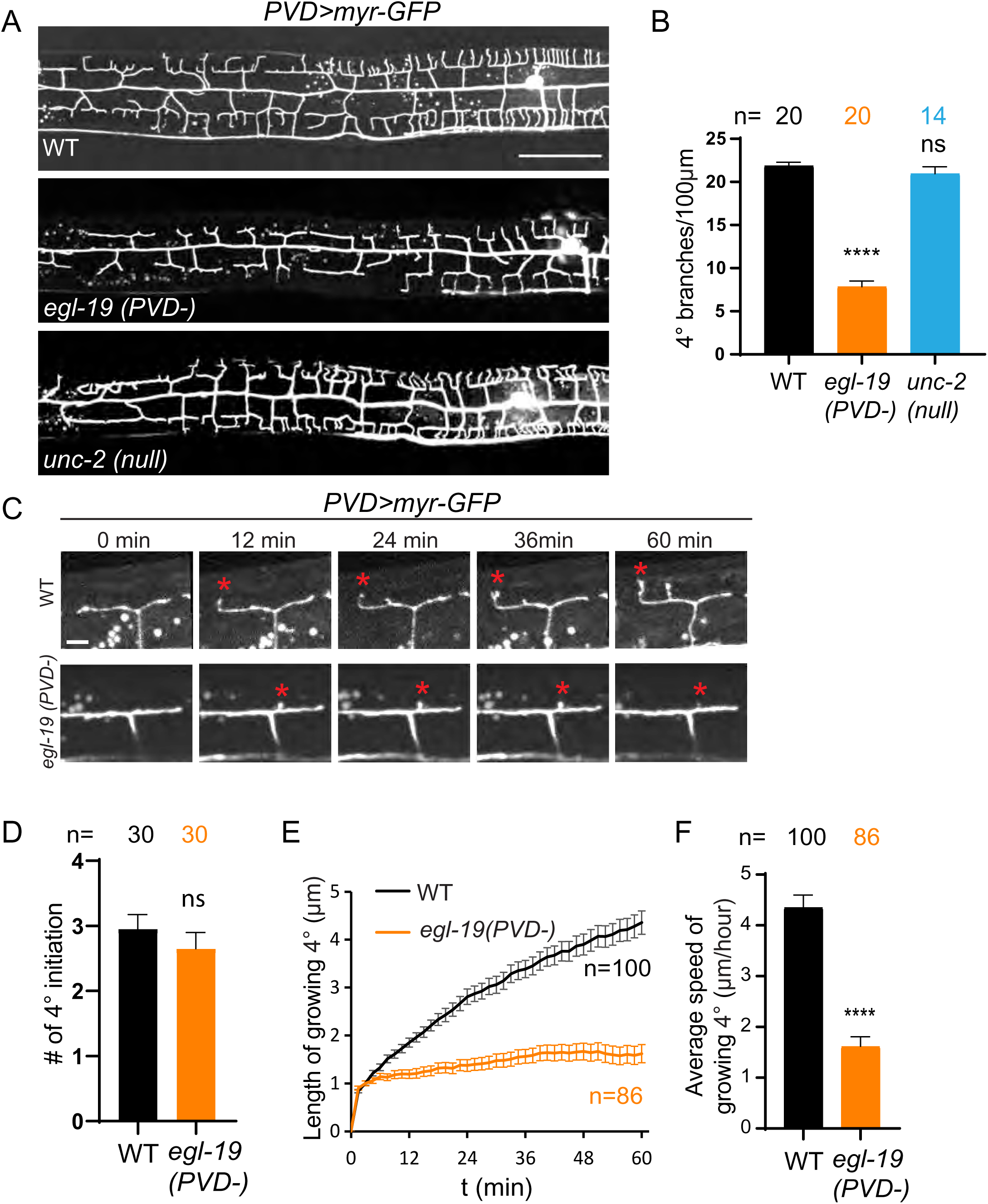
L-type VGCCs are required for dendrite morphogenesis. (**A**) Representative confocal images showing PVD morphology of WT, *egl-19(PVD-)* and *unc- 2(null)* mutants at the L4 stage. Scale bar: 50 μm. (**B**) Quantifications of the number of 4° dendrites per 100 μm in WT, *egl-19(PVD-)* and *unc-2(null)* mutants. One-way ANOVA with a Tukey correction was used for statistical analysis. ****p<0.0001. ns: not significant. Error bars: SEM. (**C**) Time-lapse imaging of growing PVD dendrites in a WT and an *egl-19 (PVD-)* animal. Scale bar: 5 μm. Red asterisk, growing 4° dendrite. (**D**) Quantification of number of 4° branches initiated from 3° in a single menorah within 1 hour. Statistics, Student’s T-test. ns: not significant. Error bars: SEM. (**E**) Plot of dendrite length over time in growing 4° dendrites in WT and *egl-19(PVD-)* animals. Error bars: SEM. (**F**) Quantification of the average speed of growing 4° dendrites in WT and *egl-19 (PVD-)* animals. Statistics, Student’s T-test. ****p<0.0001. Error bars: SEM. 10 worms were imaged for each genotype in (**D-F**).

### DEG/ENaC channels are necessary for Ca^2+^ transients and dendrite growth

In order to understand what activates L-type VGCCs during dendritic growth and whether intrinsic mechanical stimulation is involved, we searched for molecular pathways that modify both the local 4° Ca^2+^ transients and dendrite outgrowth. Specifically, we tested whether the DEG/ENaC family of mechanosensitive ion channels are involved. The DEG/ENaC family of proteins are sodium selective, non-voltage-gated channels (Eastwood and Goodman, 2012), with 30 members found in *C. elegans* genome (Hobert, 2013). At least 7 DEG/ENaC genes, including *degt-1*, *del-1, mec-10*, *unc-8, asic-1, unc-105* and *deg-1*, are expressed in PVD based on genetic and expression analyses (Huang and Chalfie, 1994; Richardson et al., 2019; Smith et al., 2010; Tao et al., 2019; Taylor et al., 2021). Deletion of individual genes including *degt-1*, *del-1, mec-10*, *unc-8, asic-1* or *unc-105* did not show an obvious dendrite outgrowth phenotype, likely due to functional redundancy (data not shown). We generated a sextuple mutant (*DEG/ENaCs Δ*) carrying deletions within 6 DEG/ENaC genes *asic-1; unc-105; unc-8; degt-1; del-1; mec-10*. *DEG/ENaCs Δ* mutants showed reduced 4° dendrites, suggesting that DEG/ENaCs are indeed required for the full dendritic arbors (Fig. 3, A, B and F).

**Figure 3.**
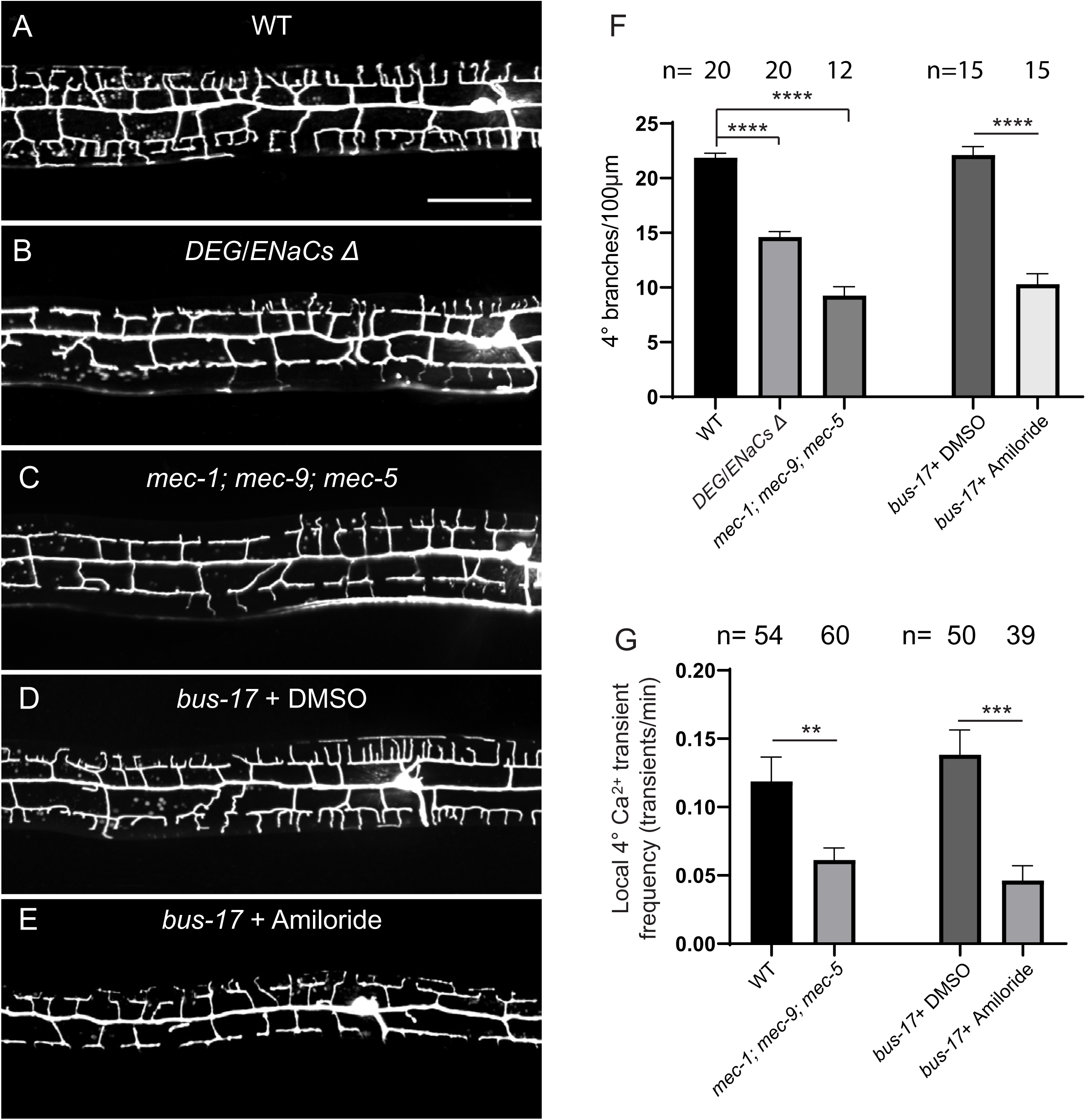
Mechanosensitive DEG/ENaC channels are required for dendritic local calcium transients and dendrite morphogenesis. (A-E) Representative confocal images of PVD morphology in the indicated genotypes. Scale bar: 50 μm. L4 animals with *PVD>myr-GFP* were imaged. *DEG/ENaCs* Δ: *asic-1; unc-105; unc-8; degt-1; del-1; mec-10.* (F) Quantification of the number of 4° dendrites per 100 μm in the indicated genotypes. Error bars: SEM. One-way ANOVA with a Tukey correction was used for comparing WT, *DEG/ENaCs* Δ and *mec-1; mec-9; mec-5*. Student’s T-test was used for comparing *bus-17* worms treated with DMSO or 3mM Amiloride. **** p<0.0001. (G) Quantification of the frequency of local 4° Ca^2+^ transients (number of events per minute) in growing 4° dendrites in the indicated genotypes. Late L3 stage were imaged. Error bars: SEM. 6 independent animals were imaged for each genotype. Statistics, Student’s T-test. ** p<0.01. *** p<0.001.

Because there might be additional DEG/ENaCs expressed in PVD, we sought to inhibit all DEG/ENaCs simultaneously by other means. First, we treated worms with Amiloride, a drug which blocks the currents of DEG/ENaC channels (Palmer, 1992). To increase the drug permeability, we performed drug treatments in *bus-17* mutants which have a thinner cuticle than wild-type (Bounoutas et al., 2009). Indeed, Amiloride treatment lowered the frequency of local 4° Ca^2+^ transients during dendrite outgrowth and reduced the number of 4° dendrites (Fig. 3, D, E, F and G). As an alternative approach to inhibit DEG/ENaC channel function globally, we deleted the extracellular matrix proteins MEC-1, MEC-5 and MEC-9, which are required for the localization and function of DEG/ENaC channels in sensory dendrites (Emtage et al., 2004). Interestingly, the *mec-1,-5,-9* triple mutants also showed dramatically reduced frequency of local 4° Ca^2+^ transients and reduced number of 4° dendrites (Fig. 3, A, C, F and G). Together, these results indicate that the DEG/ENaC family of channels is necessary for dendrite formation. Therefore, both DEG/ENaCs and Ca^2+^ influx are required for sustained dendrite outgrowth.

### Activation of mechanosensitive channels underlies developmental Ca^2+^ transients

Genetic and behavioral experiments in worms and flies suggest that DEG/ENaCs are mechanosensitive channels (Chatzigeorgiou et al., 2010; Geffeney et al., 2011; Guo et al., 2014; O’Hagan et al., 2005). To test if other mechanosensitive channels can perform similar functions during development, we overexpressed known mechanosensitive channels in the *mec-1,-5,-9* triple mutants and asked if they can rescue the defects in local 4° Ca^2+^ transients and dendrite outgrowth. We reasoned that channels that are directly gated by membrane tension might bypass the requirement of ECM molecules and should achieve some level of rescuing effects if mechanically gated signals are responsible for the Ca^2+^ influx. Specifically, we overexpressed the only *C. elegans* Piezo homolog, *pezo-1,* in PVD. Endogenous *pezo-1::gfp* is expressed at low levels in many cells (Bai et al., 2020) including PVD (fig. S2, A to B). Deletion of *pezo-1* does not cause any abnormalities in PVD dendrite growth (fig. S2, A, F to G). We used CRISPR/Cas9 to generate knock-in animals in which the *ser-2*prom3 promoter was inserted 5’ to the ATG of *pezo-1* (*PVD>pezo-1*) (fig. S2A), to drive expression of *pezo-1* only in the PVD and OLL neurons. This resulted in stable genomic overexpression of *pezo-1* in the PVD neuron, which was confirmed by inserting another GFP tag at the C terminus of *PVD>pezo-1*. *PVD>pezo-1::gfp* dramatically increased *pezo-1::gfp* intensity compared to endogenous *pezo- 1::gfp* in PVD (fig. S2, B to D) with *pezo-1::gfp* localizing along PVD dendritic branches in the *PVD>pezo-1::gfp* strain (fig. S2E).

Remarkably, overexpression of *pezo-1* in PVD (*PVD>pezo-1*) fully restored the frequency of local 4° Ca^2+^ transients in the *mec-1,-5,-9* triple mutants back to wild-type levels (Fig. 4A). Moreover, this PVD>*pezo-1* genomic overexpression also largely rescued the dendrite outgrowth defects in the *mec-1,-5,-9* triple mutants (Fig. 4, B and C). To test if mechanosensitive channels are involved in the growth of dendrites per se, we found that *mec-1,- 5,-9* triple mutants can still initiate filopodia in time-lapse recordings, but fail to extend them beyond ∼1 µm (Fig. 4, D, E and Movie S2, S4), similar to L-type VGCC knockdown (Fig. 2, C to F). PVD>*pezo-1* promoted the sustained outgrowth of dendritic filopodia in time lapse experiments (Fig. 4, D, F, G and Movie S4, S5). PVD>*pezo-1* genomic overexpression did not induce any abnormalities in PVD morphology by itself (fig. S2F). A closed-pore mutation (P1979A) in the conserved PF(X_2_)E(X_6_)W motif of *pezo-1* (Coste et al., 2015) (fig. S3A) abolished rescue (Fig. 4, B and C), suggesting that *pezo-1*’s function as an ion-channel is required to rescue the dendritic outgrowth defects of *mec-1,-5,-9* triple mutants. The expression and localization pattern of mutant PEZO-1(P1979A) in PVD was normal (fig. S3B). The *pezo- 1(P1979A)* mutant showed a reduction in brood size, similar to the *pezo-1 null* mutant (Bai et al., 2020), suggesting that the mutation effectively blocked the PEZO-1 channel (fig. S3C).

**Figure 4.**
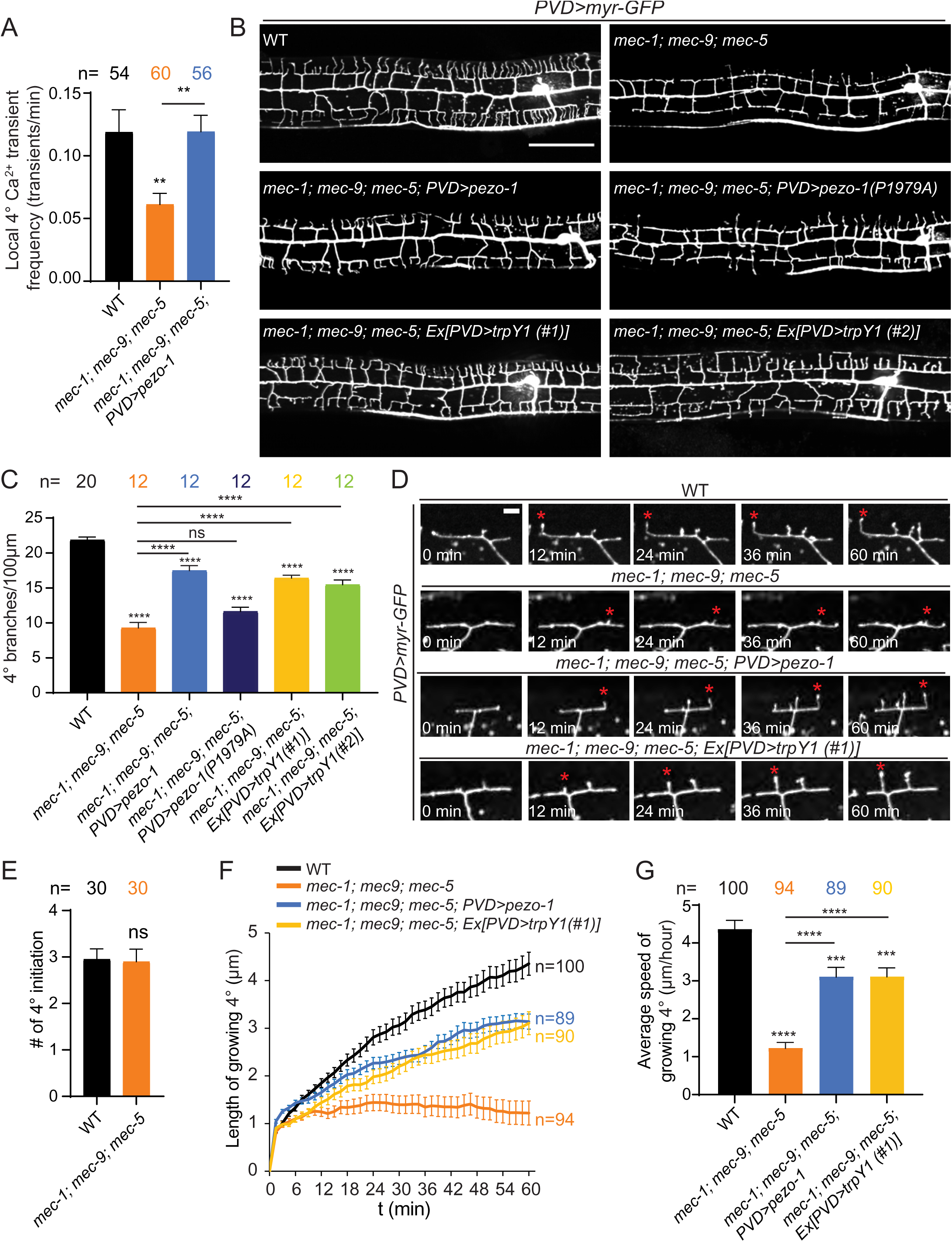
Overexpression of other mechanosensitive channels can partially rescue dendritic local calcium transients and dendrite morphogenesis defects in *mec-1,5,9* mutants. (**A**) Quantification of the frequency of local 4° Ca^2+^ transients (number of events per minute) in growing 4° dendrites in indicated genotypes at the late L3 stage. Error bars: SEM. 6 independent animals were imaged for each genotype. Statistics, One-way ANOVA with a Tukey correction. **p<0. 01. (**B-C**) Representative confocal images of PVD morphology (**B**) and quantification of the number of 4° dendrites per 100 μm (**C**) in the indicated genotypes. In (B), Scale bar: 50 μm. In (C), Error bars: SEM. Statistics, ANOVA with a Tukey correction. ****p<0.0001. ns: not significant (**D**) Time-lapse imaging of growing PVD dendrites in the indicated genotypes. Red asterisks mark the growing 4° branches. Scale bar: 5 μm. (**E**) Quantification of the number of 4° branches initiated from 3° in a single menorah within 1 hour in indicated genotypes. Error bars: SEM. Statistics, Student’s T-test. ns: not significant. (**F**) Plot showing dendrite length over time in growing 4° dendrites in the indicated genotypes. Error bars: SEM. (**G**) Quantification of the average speed of growing 4° dendrites in the genotypes shown in D and F. One-way ANOVA with a Tukey correction was used for statistical analysis. *** p<0.001, ****p<0.0001. Error bars: SEM. 10 worms were imaged for each genotype in (**E-G**).

Furthermore, transgenic overexpression of the yeast TrpY1 mechanosensitive channel was also able to partially rescue the dendrite outgrowth phenotype (Fig. 4, B to C) and partially but significantly restored the persistent 4° dendrite outgrowth in *mec-1,-5,-9* triple mutants (Fig. 4, D, F, and G and Movie S4, S6). Similar channel replacement experiments overexpressing the worm TRP-4/NOMPC failed to rescue the *mec-1,-5,-9* triple mutants (data not shown). While the Piezo and the TrpY1 channels share no sequence homology, both have been demonstrated to be activated by membrane tension and stretch without the requirement of accessory proteins (Coste et al., 2010; Coste et al., 2012; Su et al., 2011; Syeda et al., 2016; Zhou et al., 2003). In contrast, the mechanosensitive function of TRP-4/NOMPC requires microtubule tethering (Zhang et al., 2015), which is lacking during the terminal branch outgrowth of PVD (Liu et al., 2019). These pieces of evidence strongly support the notion that mechanical forces incurred at growing 4° dendrites trigger Na^+^ influx through DEG/ENaCs leading to membrane depolarization, and subsequently activate L-type VGCCs, which are required for the persistent outgrowth of dendrite branches. In wild-type animals, DEG/ENaC channels on growing dendrites sense these mechanical forces during morphogenesis to depolarize the membrane and induce Ca^2+^ influx. The ECM proteins MEC-1, -5 and -9 are required for the activation of DEG/ENaCs, and therefore are required for dendrite development. In the absence of DEG/ENaC activation in *mec-1,-5,-9* triple mutants, overexpression of PEZO-1 or TrpY1 can replace the mechanosensing function of DEG/ENaCs by sensing the morphogenesis-related membrane stretch directly. To test if DEG/ENaC channels are similarly required in other types of dendrites, we examined another neuron in *C. elegans* with highly branched dendrites, FLP. *mec-1,-5,-9* triple mutants showed reduced dendrite branching in FLP (fig. S4) suggesting this function is required in multiple neuron types.

### Overactivation of DEG/ENaC channels cause dendrite growth deficits

The genetic redundancy made it difficult for us to analyze the involvement of individual DEG/ENaCs in dendrite growth, we therefore tested the genetic interactions between DEG/ENaCs and the VGCC *egl-19* by examining gain-of-function DEG/ENaC mutations. UNC- 8 is an DEG/ENaC member that is expressed, and functions in PVD (Tao et al., 2019). The *unc- 8(e15)* allele, a gain-of-function (*gof*) allele, contains a G387E mutation which causes enhanced channel current when expressed in oocytes and led to neuronal cell death in *C. elegans* (Wang et al., 2013). *unc-8(e15)* did not result in PVD cell death, but showed reduced 4° dendrites (fig. S5, A and C). Consistent with the enhanced UNC-8 channel current, we observed a ∼400% enhancement in PVD baseline Ca^2+^ levels in *unc-8(e15)* mutants compared to wild-type (fig. S5, B and D). Both the increased baseline Ca^2+^ level and the dendrite growth phenotypes were completely restored by a weak loss-of-function *(lf)* allele of the VGCC *egl-19(ad1006)* (fig. S5, A to D). This weak allele of *egl-19* by itself did not alter PVD Ca^2+^ levels, or dendrite growth (fig. S5, C and D). *egl-19(ad1006)* mutants have previously been shown to display a normal activation time constant but smaller amplitude in Ca^2+^ recordings from body wall muscle (Gao and Zhen, 2011). This result indicates that overactivation of UNC-8 leads to excessive depolarization and L-type VGCC-dependent Ca^2+^ influx, which hampers dendrite growth.

Additionally, we identified a gain-of-function allele (*wy1014*) of the DEG/ENaC channel *del-1* in a forward genetic screen to isolate dendrite morphology mutants. *del-1(wy1014)* exhibited strong loss of PVD dendritic arbors, especially 4° branches (Fig. 5, A, B and D). DEL- 1 is also a member of the DEG/ENaC family of mechanosensitive ion channels. We have recently shown that DEL-1, UNC-8 and MEC-10 function together as sensors for proprioceptive stimuli in mature PVD neurons (Tao et al., 2019). Unlike inhibition of DEG/ENaCs in *mec-1,-5,- 9* triple mutants, time-lapse recording of outgrowth events showed that *del-1(wy1014)* mutants have both reduced 4° dendrite initiation and slower dendrite growth speed (Fig. 5, F to G and Movie S2, S7). Several lines of evidence indicate that the point mutation (P309S) found in the extracellular domain of *del-1(wy1014)* leads to a gain-of-function (*gof*) channel. First, baseline cytosolic Ca^2+^ levels are increased ∼350% in *del-1(wy1014)* mutants compared to wild-type animals (Fig. 5, C and E), like the gain-of-function *unc-8(e15)* mutant. Second, *del-1(null)* mutants had normal dendrite outgrowth (fig. S6, B to C). Third, a pore mutation completely suppressed the dendrite outgrowth defects seen in *wy1014* mutants (*wy1353* contains both the *wy1014* and the pore mutation (fig. S6, A, B and C). Fourth, a null mutation of extracellular matrix protein *mec-9*, required for DEG/ENaC channel activation, partially suppress the PVD 4° dendritic arbor defects (fig. S6, B to C). Finally, like *unc-8(e15)*, the increased Ca^2+^ levels in the *del-1(wy1014)* mutant are restored to wild-type levels in the presence of a weak loss-of-function allele of *egl-19* (Fig. 5, C and E). Importantly, while the baseline Ca^2+^ levels in the *del- 1(wy1014)* mutants were elevated, the frequency of local 4° Ca^2+^ transients were dramatically reduced (Fig. 5H), indicating that local transient increases in Ca^2+^, rather than baseline cytosolic Ca^2+^ concentration per se, are required for dendrite growth. These data suggest that overactive DEL-1 leads to excessive activation of the L-type VGCCs, which results in high levels of baseline Ca^2+^ and reduced Ca^2+^ transients during dendritic outgrowth. Consistent with this hypothesis, both DEL-1 and EGL-19 are expressed in PVD during dendrite development and localize to punctate structures along outgrowing dendritic branches (fig. S7). Together, these data indicate that the DEG/ENaC family of mechanosensitive channels activate the L-type VGCCs to generate local Ca^2+^ transients which are required for continuous dendrite outgrowth.

**Figure 5.**
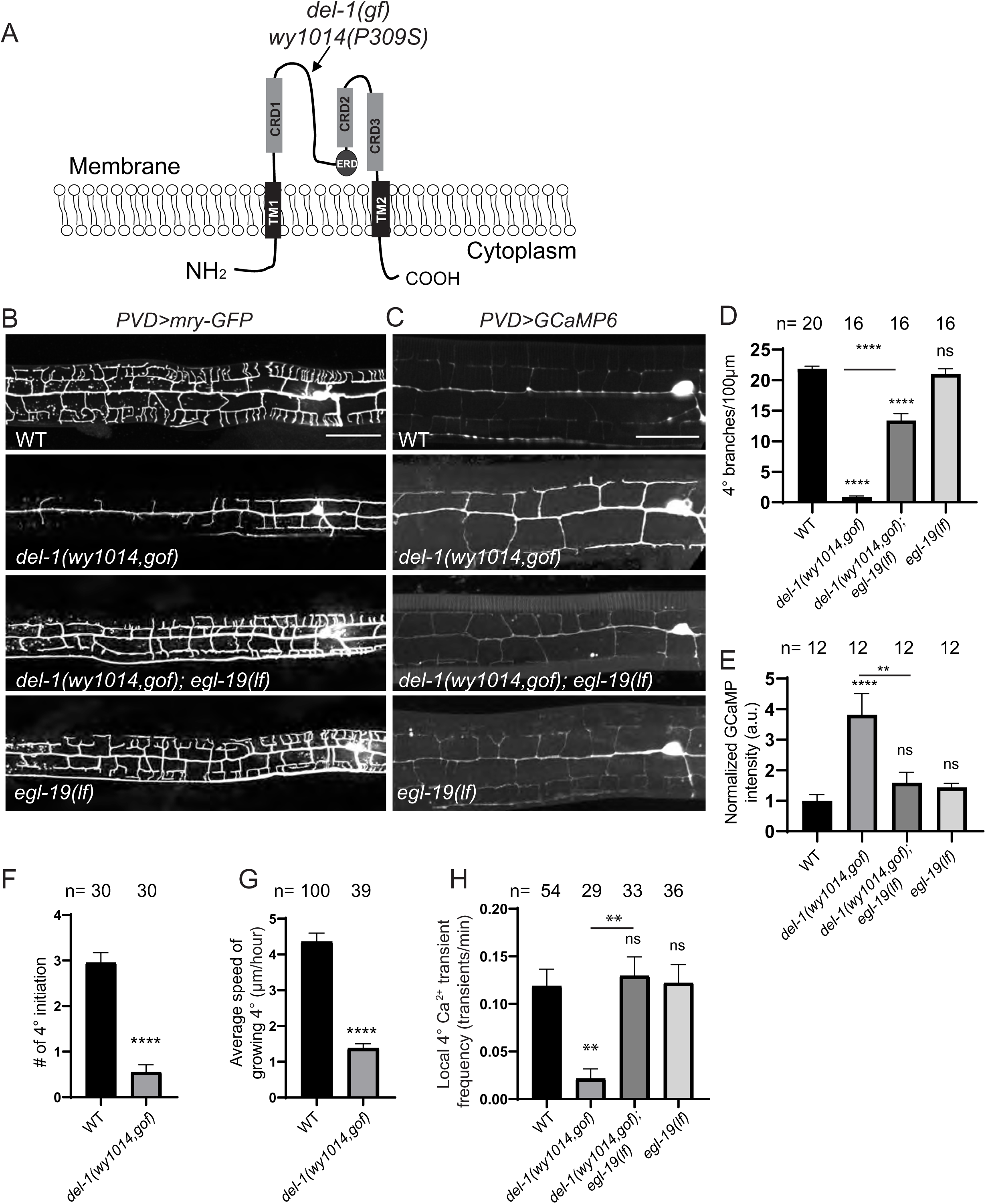
Mechanosensitive DEG/ENaC channels act upstream of the L-type VGCC *egl-19*. (**A**) Illustration of DEL-1 isoform a with the *wy1014* mutation indicated. CRD: Cysteine-rich domain. TM: transmembrane domain. ERD: Extracellular regulatory domain. (**B**) Representative confocal images of PVD morphology of indicated genotypes at the L4 stage. Scale bar: 50 μm (**C**) GCaMP6 fluorescence images in the indicated genotypes at the L4 stage. Scale bar: 25 μm. (**D**) Quantification of the number of 4° dendrites per 100 μm in the indicated genotypes. (**E**) Quantification of the mean GCaMP6 intensity within the PVD dendrite in the indicated genotypes. Normalized to WT. (**F**) Quantification of the number of 4° branches initiated from 3° in a single menorah within 1 hour in the indicated genotypes. (**G**) Quantification of the average speed of growing 4° dendrites in the indicated genotypes. In (**F-G**), Statistics, Student’s T-test. ****p<0.0001. 10 worms were imaged for each genotype. Error bars: SEM. (**H**) Quantification of local Ca^2+^ transient frequency (number of events per minute) in growing 4° dendrites in indicated genotypes at the late L3 stage. 5-6 independent animals were imaged for each genotype. In (**D**, **E**, **H**), Statistics, One-way ANOVA with a Tukey correction. **p<0. 01. ****p<0.0001. ns: not significant. Error bars: SEM.

**Figure 6.**
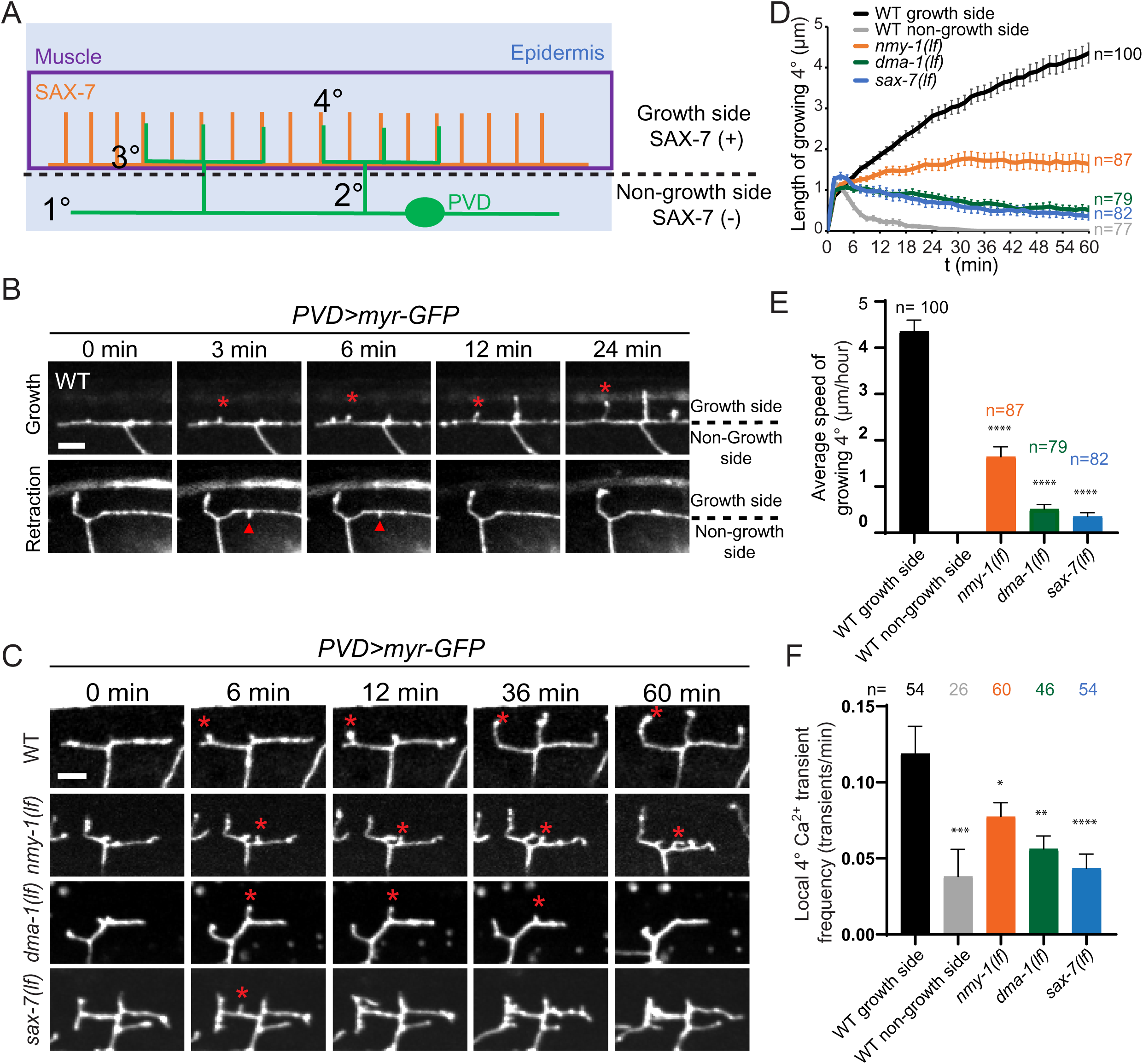
Ligand-receptor interactions and non-muscle myosin II mediate the activation of DEG/ENaC channels. (**A**) Cartoon showing the stripe-like pattern of SAX-7 (ligand) expressed in the epidermis that colocalizes with 4° dendrites. (**B**) Example time-lapse images of a 4° dendrite growing towards the growth side (upper panels, marked with asterisk) and towards the non-growth side (lower panels, marked with arrowhead). Scale bar: 5 μm. (**C**) Time-lapse images of growing 4° dendrites in the indicated genotypes. Red asterisks mark the growing 4° dendrites on the growth side. Scale bar: 5 μm. (**D**) A plot of growing 4° dendrite length over time in the indicated genotypes. Error bars: SEM. (**E**) Quantification of the average speed of 4° dendrite growth in the indicated genotypes. One-way ANOVA with a Tukey correction was used for statistical analysis. ****p<0.0001. Error bars: SEM. 10 worms per genotype were imaged. (**F**) Quantification of local Ca^2+^ transient frequency (number of events per minute) in growing 4° dendrites in indicated genotypes at the late L3 stage. Statistics, One-way ANOVA with a Tukey correction. *p<0.05, **p<0.01, ***p<0.001, ****p<0.0001. Error bars: SEM. 6 animals were imaged for each genotype.

### Guidance ligand-receptor interactions activate DEG/ENaC channels

Dendritic membranes experience multiple cellular and molecular forces during outgrowth, including adhesion with the extracellular environment and interaction with the intracellular cytoskeleton. To understand what molecular forces activate DEG/ENaCs during dendrite growth, we first tested the involvement of the dendrite adhesion ligand-receptor complex. SAX-7/L1CAM is the instructive extracellular ligand which guides PVD dendrite growth and branching by activating the transmembrane adhesion molecule DMA-1 (Dong et al., 2013; Salzberg et al., 2013). SAX-7 is present in the adjacent epidermis as stripes localized on one side of the PVD 3° dendrite, where 4° branches form (growth side), but absent from the opposite side (non-growth side) (Fig. 6A). Time-lapse imaging showed that during development, 4° dendritic filopodia initiate on both sides of the 3° branch. However, filopodia that form on the non-growth side do not extend beyond ∼1 µm in length and invariably retract (Fig. 6, B, D and E), similar to the growth dynamics of *mec-1,-5,-9* triple mutants on the growth side (Fig. 4D). Consistently, local Ca^2+^ transients within filopodia on the non-growth side are less frequent compared to those on the growth side, suggesting that ligand-receptor adhesion is required to activate local Ca^2+^ influx through DEG/ENaCs (Fig. 6F).

To test the causality of ligand-receptor interactions and local 4° Ca^2+^ transients, we analyzed the 4° dendrite growth phenotypes in weak loss-of-function *(lf)* alleles of *sax-7* and *dma-1*. We chose to use weak loss-of-function alleles because *sax-7* and *dma-1* null mutations cause completely disordered and truncated dendrites with no 4° dendrite growth. Weak loss-of-function alleles of *sax-7* and *dma-1* showed mild defects in 3° branches, but dramatically reduced 4° branches (fig. S8, A and B). Indeed, partial loss of SAX-7 or DMA-1 caused reduced 4° Ca^2+^ transients in the filopodia on the growth side and a severely reduced outgrowth speed of 4° dendrites (Fig. 6, C to F and Movie S8, S9). These results indicate that ligand-receptor interactions contribute to the mechanical force that activates DEG/ENaCs. Next, we examined the non-muscle myosin *nmy-1* partial loss-of-function mutant which showed slightly fewer, and significantly shortened 4° dendrites (fig. S8). We found that it also displayed a reduced frequency of local 4° Ca^2+^ transients as well as a reduced speed of 4° dendrite growth, indicating that myosin contractility also contributes to DEG/ENaC activation (Fig. 6, C to F and Movie S10).

Together, these data indicate that the initial filopodia formation of 4° dendrites is random in growth direction. Appropriate ligand-receptor interactions, and actin-myosin contractility, generate mechanical forces on the “growth side” which activate DEG/ENaCs, depolarizing the membrane to trigger local 4° Ca^2+^ transients. Since the ligand SAX-7 does not localize to the non-growth side, filopodial growth on this side does not trigger DEG/ENaC mediated Ca^2+^ transients. Excessive, or insufficient activation of DEG/ENaCs leads to reduced local 4° Ca^2+^ transients and truncated dendrite outgrowth.

To further test this model, we investigated genetic interactions between the gain-of-function DEG/ENaC mutation and mutants with defects in adhesion, or myosin contractility. We reasoned that partial loss of adhesion, or myosin contractility, might reduce the overactivation of mutant DEG/ENaCs back to wild-type levels and correct the excessive Ca^2+^ influx caused by the overactive *del-1(gof)* mutation, *wy1014*. The *del-1(wy1014)* allele can be partially suppressed by *mec-9*, suggesting that the mutated channel is not constitutively open and can still be gated by ECM (fig. S6, B and C). Indeed, the weak allele of *sax-7* and loss-of-function in *nmy-1* significantly reduced the baseline Ca^2+^ in PVD back to wild-type levels in *del-1(wy1014)* mutants (Fig. 7, A and C) and increased PVD dendrite branch number (Fig. 7, B and D), indicating that the adhesion molecule SAX-7 and non-muscle myosin NMY-1 are required for the activation of the DEL-1 channel.

**Fig. 7.**
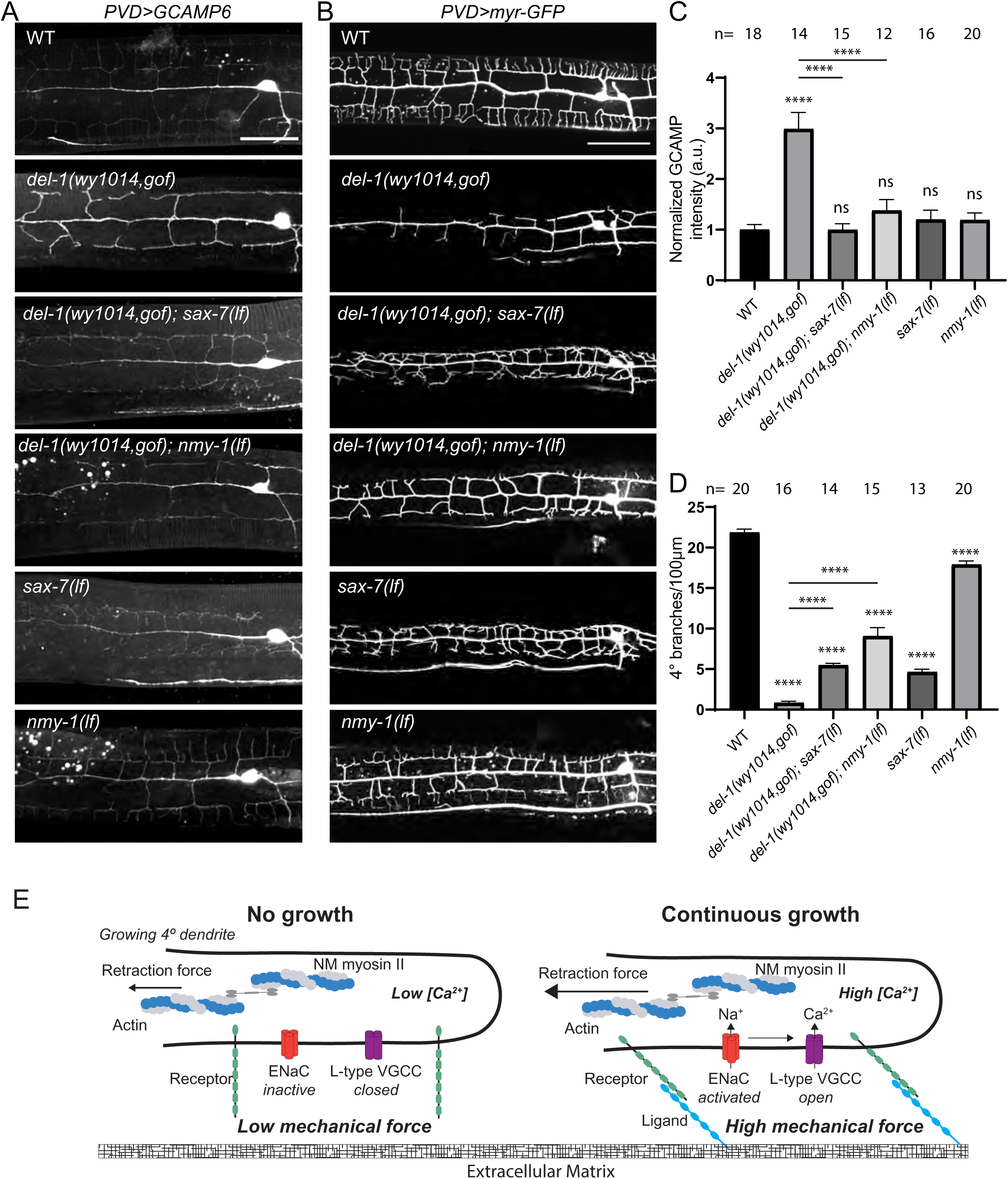
The increased Ca^2+^ levels and PVD morphology defects observed in *del-1 (wy1014)* mutants require both *sax-7* and *nmy-1*. (A) GCaMP6 fluorescence images in the indicated genotypes of L4 stage worms. Scale bar: 25μm. (**B**) Confocal images showing PVD morphology of indicated genotypes at the L4 stage. Scale bar: 50μm. (**C**) Quantification of the mean GCaMP6 intensity within the PVD dendrite in the indicated genotypes (normalized to WT). (**D**) Quantification of the number of 4° dendrites per 100 μm in the indicated genotypes. In (**C** and **D**), ANOVA with a Tukey correction was used for statistical analysis. ****p<0.0001. ns. not significant. Error bars: SEM. (**E**) A cartoon showing the proofreading mechanism during dendrite outgrowth.

## Discussion

In summary, we show that mechanosensitive channels function as part of a proofreading mechanism to regulate dendrite outgrowth. During development, dynamic filopodial growth and retraction sculpt the shape of dendritic arbors. During growth, filopodia experience mechanical force generated by correct ligand-receptor interactions which activate mechanosensitive DEG/ENaCs, triggering transient local Ca^2+^ influx through L-type VGCCs. These local Ca^2+^ transients further stimulate the continuous growth of dendritic branches, without which filopodia are retracted (Fig. 7E).

Both spontaneous and evoked activity during development shapes neural circuits (Pan and Monje, 2020). The nature of the spontaneous activity related to cellular morphogenesis has been elusive. Ligand independent intrinsic activity of olfactory receptors were postulated to underlie wiring decisions in the olfactory system (Imai and Sakano, 2008). This mechanism is likely specific to the olfactory neurons due to the specific expression of receptor genes in the olfactory sensory neurons. A number of studies have shown that Ca^2+^ transients can influence guidance decisions by modulating growth cone cytoskeleton (Gasperini et al., 2017). However, what *in vivo* stimuli triggers Ca^2+^ transients in the developing axon is largely unknown. Here we show that the vast majority of local Ca^2+^ transients in dendrites are associated dendrite outgrowth. First, many Ca^2+^ transients appear to be restricted to the tip of dendritic filopodia, the loci of active growth. Second, the frequency of local Ca^2+^ transients is higher during dendrite outgrowth compared to mature arbors. Third, local Ca^2+^ transients require correct ligand-receptor interactions, which pattern dendrite outgrowth, and non-muscle myosin. Given that many neurite guidance and synaptogenesis receptors are cell adhesion molecules, it is conceivable that interactions between guidance molecules together with non-muscle myosin mediated contractility generate molecular traction force, which is then transduced via mechanosensitive ion channels into localized Ca^2+^ transients capable of modulating growth cone behavior.

In PVD, the DEG/ENaC mechanosensitive channels are responsible for activating the L- type VGCCs during dendrite growth. Our genetic data revealed two surprising aspects of DEG/ENaC involvement. First, there appears to be a great deal of redundancy among DEG/ENaC subunits in responding to neurite growth related force. While global inhibition of DEG/ENaCs and gain-of-function mutations in *del-1* and *unc-8* caused dramatic growth defects, loss of single DEG/ENaC genes did show dendrite growth phenotypes. Second, overexpression of the *C. elegans* Piezo, *pezo-1*, or the yeast stretch activated channel TrpY1, was sufficient to bypass the function of DEG/ENaCs and support dendrite growth. DEG/ENaCs, Piezo channels and TrpY1 have different gating mechanisms and generate diverse currents (Coste et al., 2010; O’Hagan et al., 2005; Palmer et al., 2001) Given that all three can support dendrite growth, it is likely that the precise nature of the mechanosensitive response might not be critical. Instead, the general mechanosensitive property of the developing neurite appears to trigger depolarization and induce local Ca^2+^ transients. Mechanosensitive ion channels are widely expressed in the nervous system and can potentially be a general mechanism during the neuronal morphogenesis process.

Guidance receptors and ligands instruct axons and dendrites to form stereotyped projections and branches *in vivo*. How do the mechanosensitive signals intersect with the classic axon guidance receptor and ligands? We found that the local Ca^2+^ transients induced by mechanical stimuli were dependent on the same ligand-receptor interactions required for dendrite guidance. Furthermore, genetic interaction analyses showed that adhesion between ligand and receptors functioned upstream of DEG/ENaC channels. Together, these data are consistent with a model in which the mechanosensitive ion channels function as a “molecular proprioceptor” to proofread the correct guidance interactions. In this model, the initial filopodia formation is random. If a filopodia contacts the correct ligand, it will allow ligand receptor binding to occur. The adhesiveness, together with actin-myosin contraction, generates traction force and activates the mechanosensitive channels to generate local Ca^2+^ transients. Filopodia with Ca^2+^ transients continue to elongate and branch while those without Ca^2+^ transients will retract. A number of cytoskeletal modulators can be regulated by Ca^2+^ (Gasperini et al., 2017). Future studies will shed light on the molecular events that are modulated by Ca^2+^ and whether this is a general mechanism for neuronal morphogenesis.

## Acknowledgments

We thank Liqun Luo for critical reading of the manuscript and helpful suggestions; Roderick MacKinnon for advice regarding trpY1 rescue experiment; Callista Yee for providing a list of PVD expression genes; members of the Shen lab for sharing strains and discussions. We thank Massimo Hilliard for his generosity and insightful discussions. We thank Yuhan Zhang for technical assistance and CGC for strains. This work was funded by the Howard Hughes Medical Institute, of which K.S. is an investigator. S.C received funding from NHMRC-ARC Dementia Development Fellowship APP1108489.

## Author contributions

Conceptualization: K.S., L.T., S.C.

Investigation: L.T., S.C.

Visualization: L.T., S.C.

Funding acquisition: K.S.

Project administration: K.S.

Resources: R.S

Supervision: K.S.

Writing – original draft: K.S., L.T.

Writing – review & editing: S.C., R.S.

## METHODS

### Strains

*C. elegans* strains were grown on nematode growth medium (NGM) plates seeded with OP50 *E. coli.* at 20°C or room temperature (22.5°C), except for the strains containing the temperature sensitive allele *wy50544* that were grown at 16°C, imaged at room temperature. N2 Bristol was used as the wild-type strain. All the strains used are listed in Table S1.

### Plasmids and transgenes

All the plasmids used for *C. elegans* expression were generated using the pSM_delta_ vector backbone, a derivative of pPD49.26 (A. Fire). Transgenic worms were made using standard gonad transformation by injection of the plasmid with co-injection marker (Mello and Fire, 1995). trpY1 cDNA was codon optimized and synthesized in IDT (Integrated DNA Technologies)*. Pser2prom3* (4138bp) is used for PVD expression. *Pmec-3* is used for FLP expression.

### CRISPR/Cas9-mediated genome editing

Endogenous insertions, point mutations and deletions were all generated by gonadal microinjection. 1.52μM cas9 protein, 1.52μM tracer RNA, 1.52μM guide RNA, and either 0.5 μM single-stranded DNA (for point mutation or small insertion) or 0.2-2μM PCR products (for GFP or GFP FLPon insertions, *Pser2prom3*) as repair templates mixed with co-injection marker (rol-6) were injected into the worms. Two guide RNA and no repair template were used to generate deletions. ssDNA repair templates were designed with 50-75 bp homology sequence. PCR products were amplified from plasmid containing *Pser2prom3*, GFP, AID-ZF-GFP or from pSK–FLPon-GFP vectors with ultramer oligonucleotides containing ∼100bp homology sequence. The guide RNAs and repair templates for each gene are listed in Table S2. A short promoter of *ser2prom3* (1618bp) was inserted into the endogenous pezo-1 promoter region. FLP-on expression of fluorescent tags (Schwartz and Jorgensen, 2016) was used for PVD cell-specific, endogenous GFP knock-in.

### Calcium imaging during PVD dendrite development

Animals were cultured on nematode growth medium (NGM) plates seeded with *E. coli* OP50 and imaged at late L3 or 1-day old adults. Prior to imaging animals were placed on an unseeded NGM plate and washed in a ∼20 µL drop of M9 buffer to remove excess *E. coli*. ∼3-4 animals were transferred to a 1.5 µL drop of 10 mM levamisole on a 30 mm glass-bottom petri dish (MatTek). 5% agarose pads were made on 15 mm circular glass coverslips (Fisherbrand) and inverted onto the dish containing immobilized animals.

For *bus-17* worms, late L3 stage *bus-17* worms were picked on the seeded NGM plate with 3mM amiloride or control plate (DMSO) for 2 hours treatment. then worms were mounted onto a glass-bottom petri dish as described above. 0.25mM levamisole and 5% agarose pads containing DMSO (control) or amiloride (3mM) were used.

Animals were imaged continuously for 1 hour with a 5 sec frame rate (z-stack) or 15-30 min with a 1 sec frame rate on a CSU-X1 spinning disk (Yokogawa), QuantEM:512SC Hamamatsu camera, 40x objective, using excitation wavelengths of 488 nm and 561 nm.

### Dynamic imaging of dendrite development

The late L3 stage worms were immobilized on 5% agar pad using 10 mM levamisole in a glass-bottom petri dish as above. Imaging was performed in a spinning disk confocal microscope with CSU-X1 spinning disk (Yokogawa), QuantEM:512SC Hamamatsu camera, a 40× objective, 488 nm nm laser. 2- hour movies were recorded with 90 sec intervals between acquisitions, z-stacks.

### Confocal imaging

L4 stage hermaphrodite worms were immobilized on 3.5% agar pads using 10 mM levamisole in M9 buffer before imaging. To examine PVD morphology, PVD>GCaMP6 level, *PVD>pezo- 1::GFP* and *pezo-1-FLPpon::GFP*, imaging were performed on a spinning disk system (3i) with a CSU-W1 spinning disk (Yokogawa), 488-nm lasers, 40x/0.9 or 63×/1.2 NA water-immersion objective, and a Prime95B camera (Photometrics).

To look at the subcellular localization of EGL-19::GFP, PEZO-1::GFP, images were acquired using a spinning disk confocal microscope with QuantEM:512SC Hamamatsu camera, a 63×/1.4 NA or 100x/1.4 NA objective, 488 nm and 561 nm lasers. Z-stacks were collected.

### Amiloride treatment

Amiloride (Sigma) was dissolved in DMSO to a stock concentration of 300mM and then the stock was added into NMG as a final concentration of 3mM. the same amount of DMSO was added into the media as a control. To examine the PVD morphology, *bus-17* worms with PVD marker were transferred onto the seeded NGM plate with 3mM amiloride or control plate (DMSO) at the L2 stage. After 16–18-hour treatment, worms were picked for imaging at L4 stage.

### Brood size assay

Single mid-L4 hermaphrodites were picked on NGM plates seeded with OP50 and left for 24 hours to allow them to lay eggs. Then the same hermaphrodite was picked to a new seeded NGM plate for another 24h, then removed from the plate. The hatched L1 worms were counted as brood size. 10 L4 worms were used for each genotype.

### Image analysis and quantification

Images were processed and analyzed using ImageJ/Fiji (National Institutes of Health). All the images were maximum projected, except for when indicated otherwise. PVD morphology and GCaMP6 expression images were straightened, cropped and assembled into Illustrator. All statistical calculation and graphs were generated using Prism 8. The details of statistics analysis are presented in the figure legends.

For quantification of local calcium transient frequency, raw images were maximum intensity projected, and registered using the MultiStackReg plugin on ImageJ (Thevenaz et al., 1998). For each growth event, if the 4° dendrite was already initiated from the 3° at the beginning of the movie, all the frames available were used for quantification. If the 4° dendrite was not initiated from 3° at the beginning of the movie, the frame immediately before the 4° dendrite was observed protruding from the 3° branch was defined as time 0 min. And the frames from -1 min (12 frames before 0 min) to the end of the movie were used for quantification. For the non-growth events (“static”), all the frames available were used for quantification. Regions of interest (ROIs) were drawn around growing 4° dendrite branches to measure GCaMP6 intensity and in regions adjacent to growing dendrites for background subtraction. GCaMP6 intensities for each ROI were corrected for background and extracted from timelapse videos. GCaMP6 intensity traces were corrected for time-course photobleaching, and the first 10 frames were used to establish a baseline fluorescence (F) for ΔF/F calculations. For the quantification of 1 sec and 5 sec frame-rate imaging, traces were smoothed using a three-time point rolling average and peaks with amplitudes greater than 0.5 *ΔF/F* were identified. Peak counts were manually confirmed by visual inspection of the timelapse videos. Traces illustrating the calcium dynamics of 1 sec frame-rate imaging were processed as above, but not smoothed prior to display. 1 sec frame-rate imaging were used in Fig. 1B-C, and fig. S1. 5 sec frame-rate imaging were used for all the other quantifications. All the quantification in the Figures shown are measurement of calcium transients in growing 4° dendrites in the growth side, except in Figure 6 F, the non-growth side events in WT were also measured.

For quantification of dendrite growth speed, 6-12 4° dendrites in the growth side or non-growth side were randomly picked in each worm. Length of individual dendrites were tracked for 1 hour. The frame immediately before the dendrite was observed protruding from the 3° branch was defined as time 0 min. All the data shown in the Figures are 4° dendrites growth events in the growth side, except in Figure 6 D and E, the non-growth side events in WT were also quantified. 10 worms for each genotype were used for quantification. Statistical calculation and graphing were done using Prism 8.

For quantification of 4° dendrite initiation, all the protrusions from a 3° branch in one menorah were counted through the entire movie. The data was presented as number of protrusions per hour. Three menorahs were randomly picked from each worm, and 10 worms were quantified for each genotype.

For quantification of number of PVD 4° dendrite, images acquired from one worm were stitched together using ImageJ. All the branches were counted for each worm and normalized by the length of 1° dendrite (number of 4° dendrite/100um).

For quantification of GCaMP6 fluorescent intensity in the PVD dendrite, 3 lines were drawn along each type of dendrite (1°, 2°, 3° and 4°), and adjacent regions were used as the background. The mean intensity of all dendrite ROIs was calculated. Data was normalized to wild-type.

## Supplemental Movie titles

**Movie S1.** Example of 1 sec frame rate movie of local 4° Ca^2+^ Transients during PVD neuron development. GCaMP6 (green) and mCherry (magenta), WT. scale bar: 5 μm. Arrowhead: growing 4° dendrite.

**Movie S2.** Development of terminal 4° dendritic branches in a WT animal with *PVD> myr- GFP.* Scale bar: 10 μm. Arrowhead: growing 4° dendrite.

**Movie S3.** Development of terminal 4° dendritic branches in an *egl-19 (PVD-)* animal with *PVD> myr-GFP*. Scale bar: 10 μm. Arrowhead: growing 4° dendrite.

**Movie S4**. Development of terminal 4° dendritic branches in a *mec-1; mec-9; mec-5* animal with *PVD> myr-GFP*. Scale bar: 10 μm. Arrowhead: growing 4° dendrite.

**Movie S5.** Development of terminal 4° dendritic branches in a *mec-1; mec-9; mec-5; PVD>pezo-1* animal with *PVD> myr-GFP*. Scale bar: 10 μm. Arrowhead: growing 4° dendrite.

**Movie S6.** Development of terminal 4° dendritic branches in a *mec-1; mec-9; mec-5; Ex[PVD>trpY1 (#1)]* animal with *PVD> myr-GFP*. Scale bar: 10 μm. Arrowhead: growing 4° dendrite.

**Movie S7.** Development of terminal 4° dendritic branches in a *del-1(wy1014, gof)* animal with *PVD> myr-GFP*. Scale bar: 10 μm. Arrowhead: growing 4° dendrite.

**Movie S8.** Development of terminal 4° dendritic branches in a *dma-1(lf)* anima with *PVD> myr- GFP*. Scale bar: 10 μm. Arrowhead: growing 4° dendrite.

**Movie S9.** Development of terminal 4° dendritic branches in a *sax-7(lf)* animal with *PVD> myr- GFP*. Scale bar: 10 μm. Arrowhead: growing 4° dendrite.

**Movie S10.** Development of terminal 4° dendritic branches in a *nmy-1(lf)* animal with *PVD> myr-GFP*. Scale bar: 10 μm. Arrowhead: growing 4° dendrite.

**Supplementary tables**

## Supplemental figure titles and legends

**Fig. S1.**
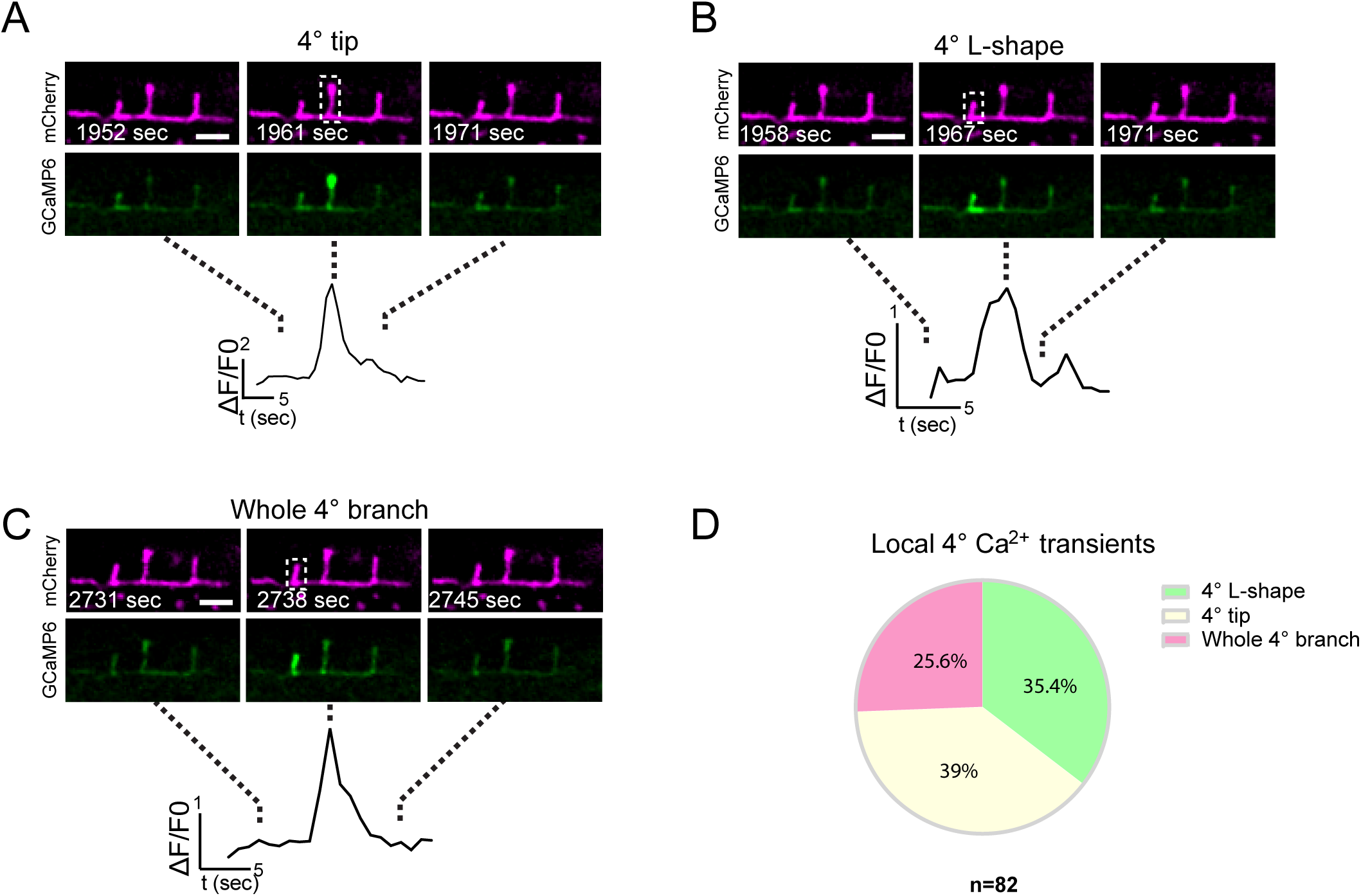
Examples of local 4° Ca^2+^ transients in developing dendrites, related to Figure 1. (A-C) Fluorescence images of GCaMP6 (green) and mCherry (magenta), and corresponding Ca^2+^ traces, showing Ca^2+^ increase in the tips of 4° dendrites (A), extending from 4° to part of the 3°dendrite (L-shape) (B), and along the whole 4° dendrite (C). Scale bar: 5 μm. (D) The percentage of each type of 4° local Ca^2+^ transient observed. Rectangles: Regions for GCaMP6 intensity measurement.

**Fig. S2.**
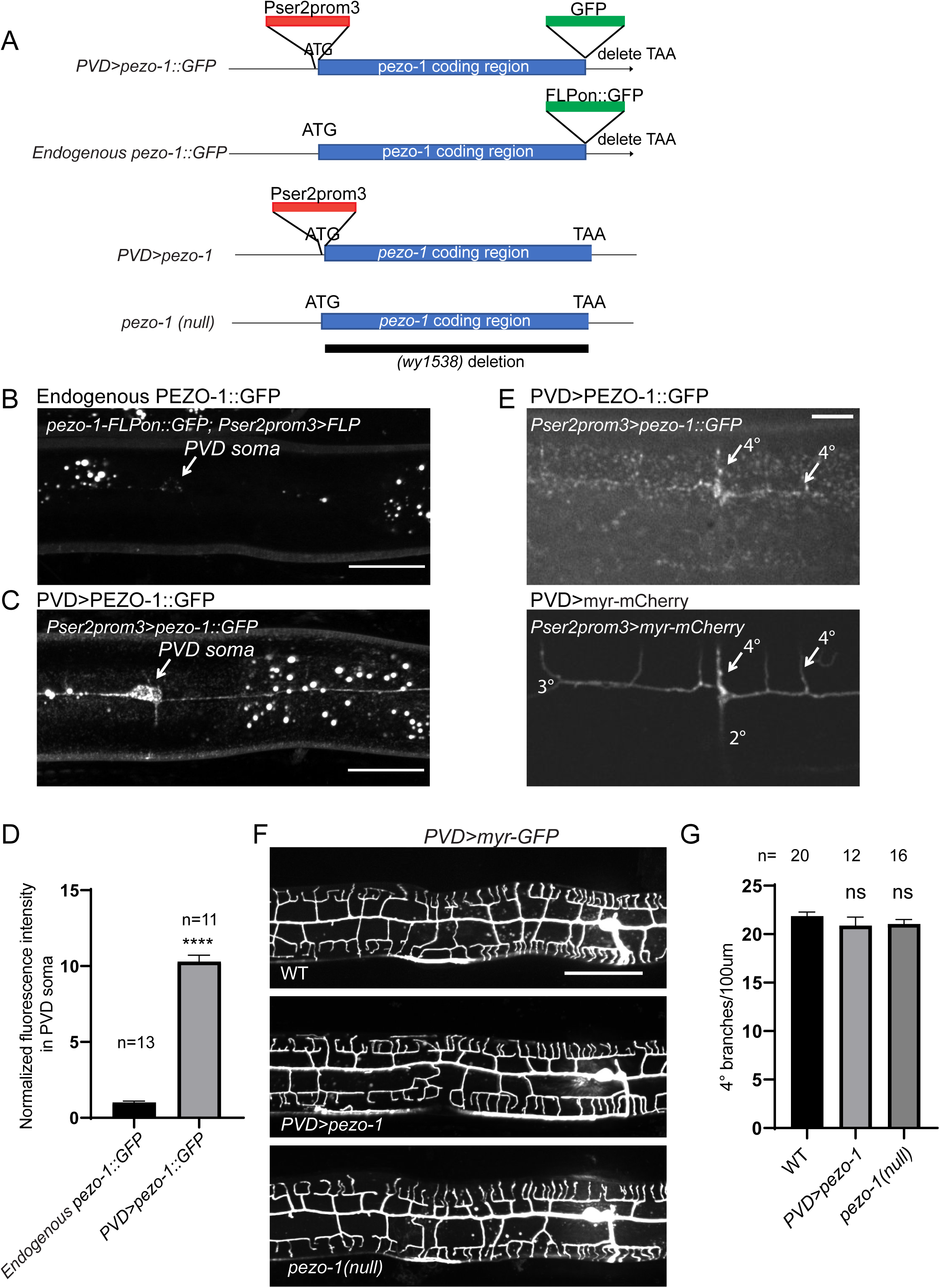
CRISPR/Cas9 insertion of the Pser2prom3 sequence into the promoter region of *pezo-1* causes overexpression of PEZO-1 in PVD, related to Figure 4. (A) *pezo-1* alleles generated with CRISPR/Cas9. (B-C) Representative confocal images of PVD cell-specific, endogenous tagged *pezo-1::GFP* (B) and *PVD>pezo-1::GFP* (C). Scale bar: 25 μm. (D) Quantification of fluorescence intensity of PEZO-1 in PVD soma, normalized to endogenous GFP-tagged pezo-1. Student’s T-test was used for statistical analysis. ****p<0.0001. Error bars: SEM. (E) Dendritic localization of PEZO-1 in *PVD>pezo-1::GFP* worms. Scale bar: 5 μm. (F) Representative confocal images of PVD morphology in the indicated genotypes. Scale bar: 50 μm. (G) Quantification of the number of 4° dendrites per 100 μm in WT, *PVD>pezo-1* and *pezo-1(null)* animals. One-way ANOVA with a Tukey correction was used for statistical analysis. ns: not significant. Error bars: SEM.

**Fig. S3.**
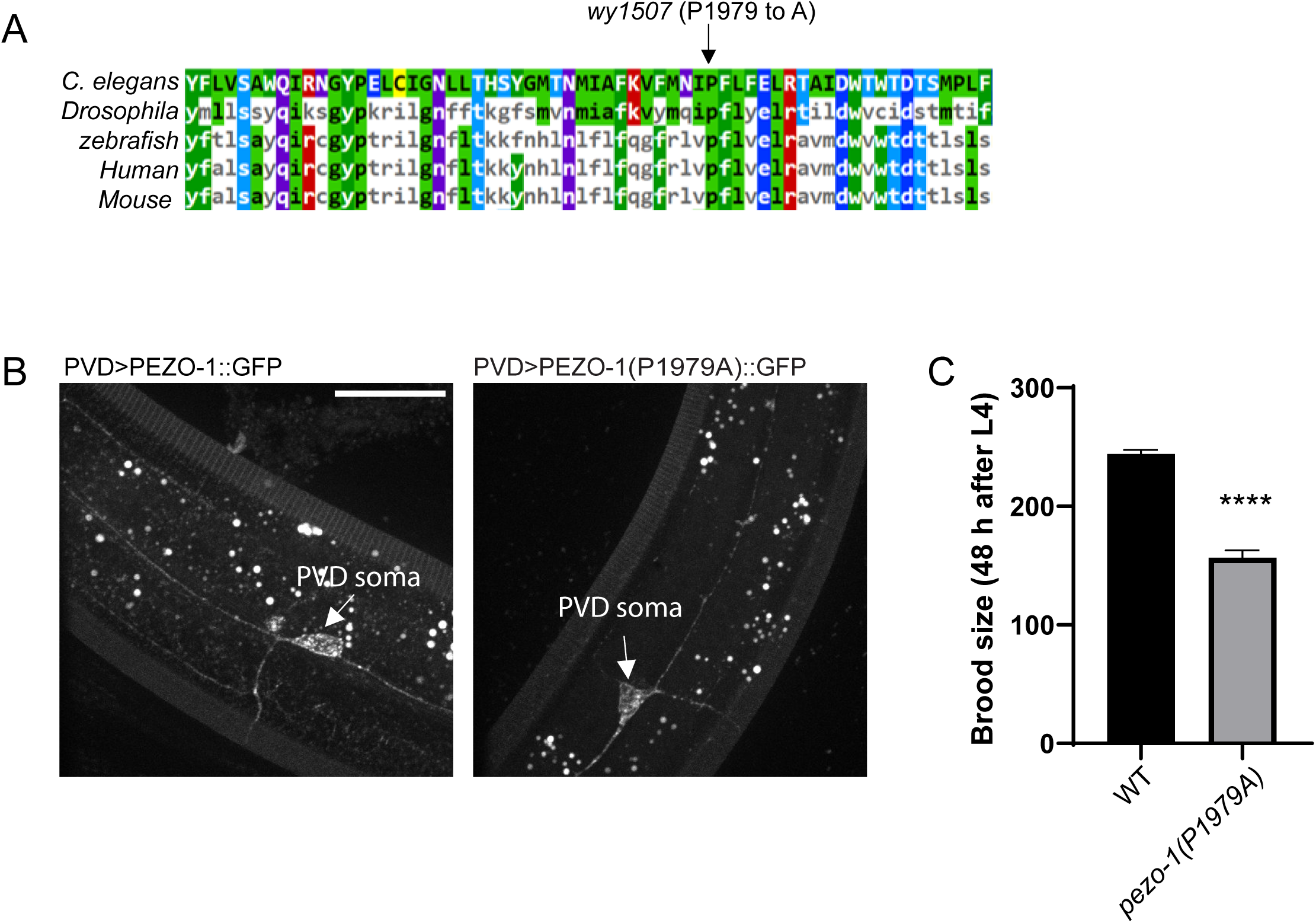
The P1979 to A mutation in PEZO-1 inhibits channel activity but shows normal localization, related to Figure 4. (A) Sequence alignment of mouse, human, fly, zebrafish, and worm Piezo-1 channels showing the conserved PF(X_2_)E(X_6_)W motif. The *wy1507* allele contains point mutations that causes a P1979 to A amino acid change. (B) Representative confocal images of both *PVD>pezo-1::GFP* and *PVD>pezo-1(P1979A)::GFP* worms. Scale bar: 25μm. (C) Brood size of WT (n=10) and *pezo-1(wy1507) (*n=10). Student’s T-test. ****p<0.0001. Error bars: SEM.

**Fig. S4.**
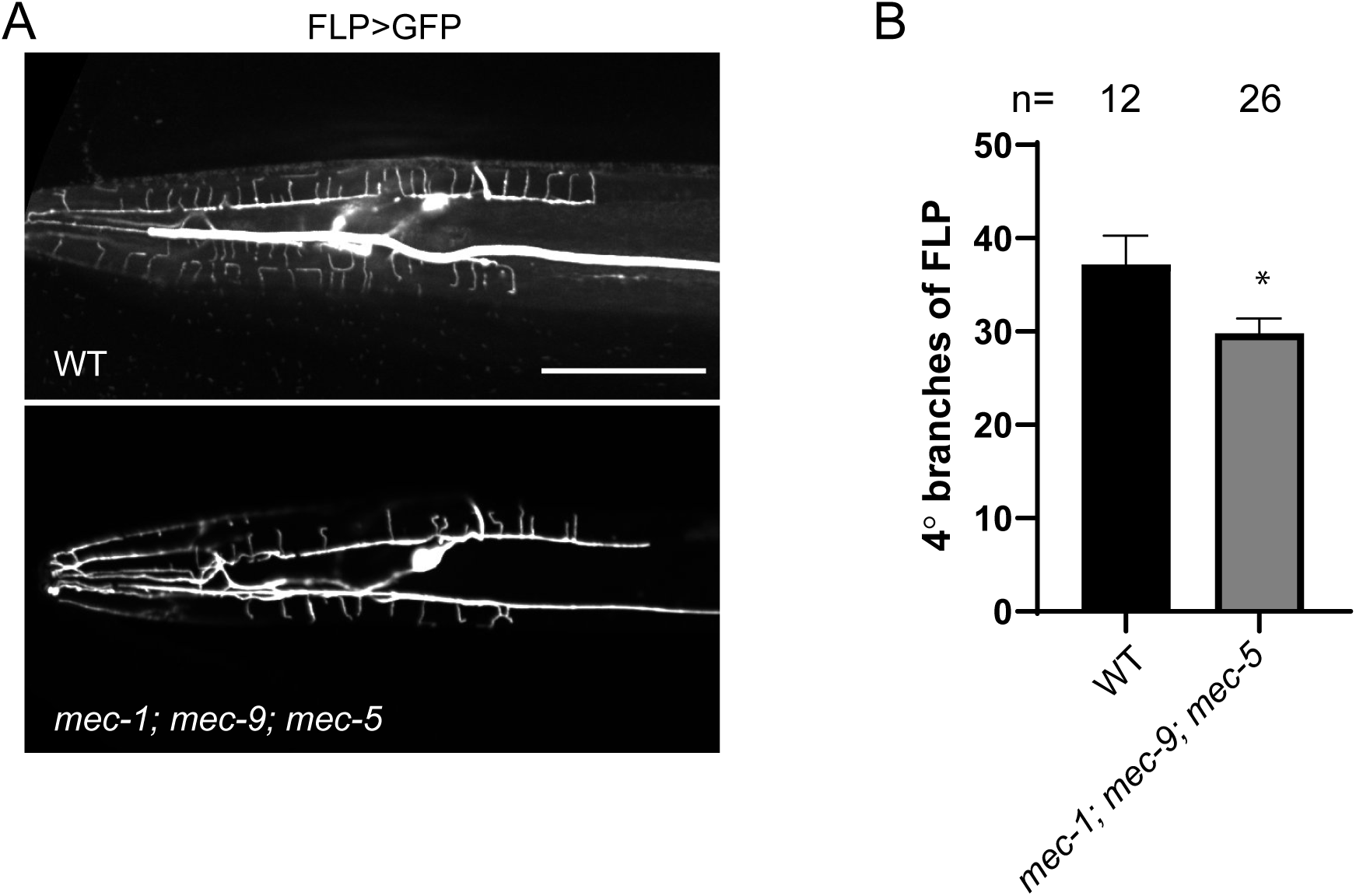
Mechanosensitive DEG/ENaC channels regulate FLP neuron morphology, related to Figure 4. (A) Confocal images showing FLP morphology of indicated genotypes in the L4 stage. Scale bar: 50μm. (B) Quantification of the number of 4° dendrites in the indicated genotypes. *p<0.05. Student’s T-test. Error bars: SEM.

**Fig. S5.**
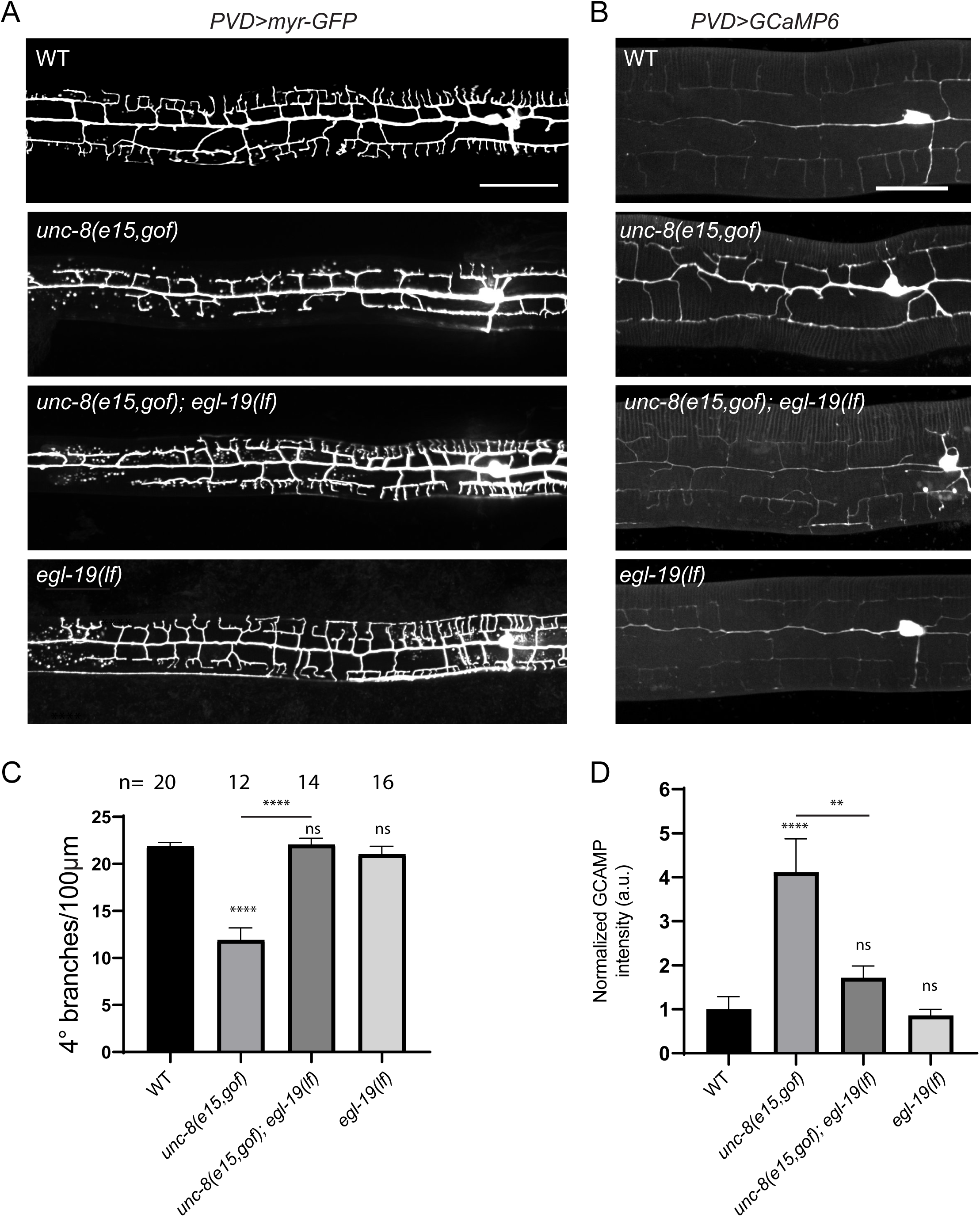
An *unc-8* gain of function allele (*e15)* shows both PVD morphology defects and increased Ca^2+^ levels that are *egl-19* dependent, related to Figure 5. (A) Confocal images showing PVD morphology of indicated genotypes in the L4 stage. Scale bar: 50 μm. (B) GCaMP6 fluorescence images in the indicated genotypes of L4 stage worms. Scale bar: 25 μm (C-D) Quantifications of the number of 4° dendrites per 100 μm (C) and mean dendritic GCaMP6 intensity (normalized to WT) in the indicated genotypes (D). One-way ANOVA with a Tukey correction was used for statistical analysis. ****p<0.0001. ns: not significant. Error bars: SEM. n=10 for each genotype in (D).

**Fig. S6.**
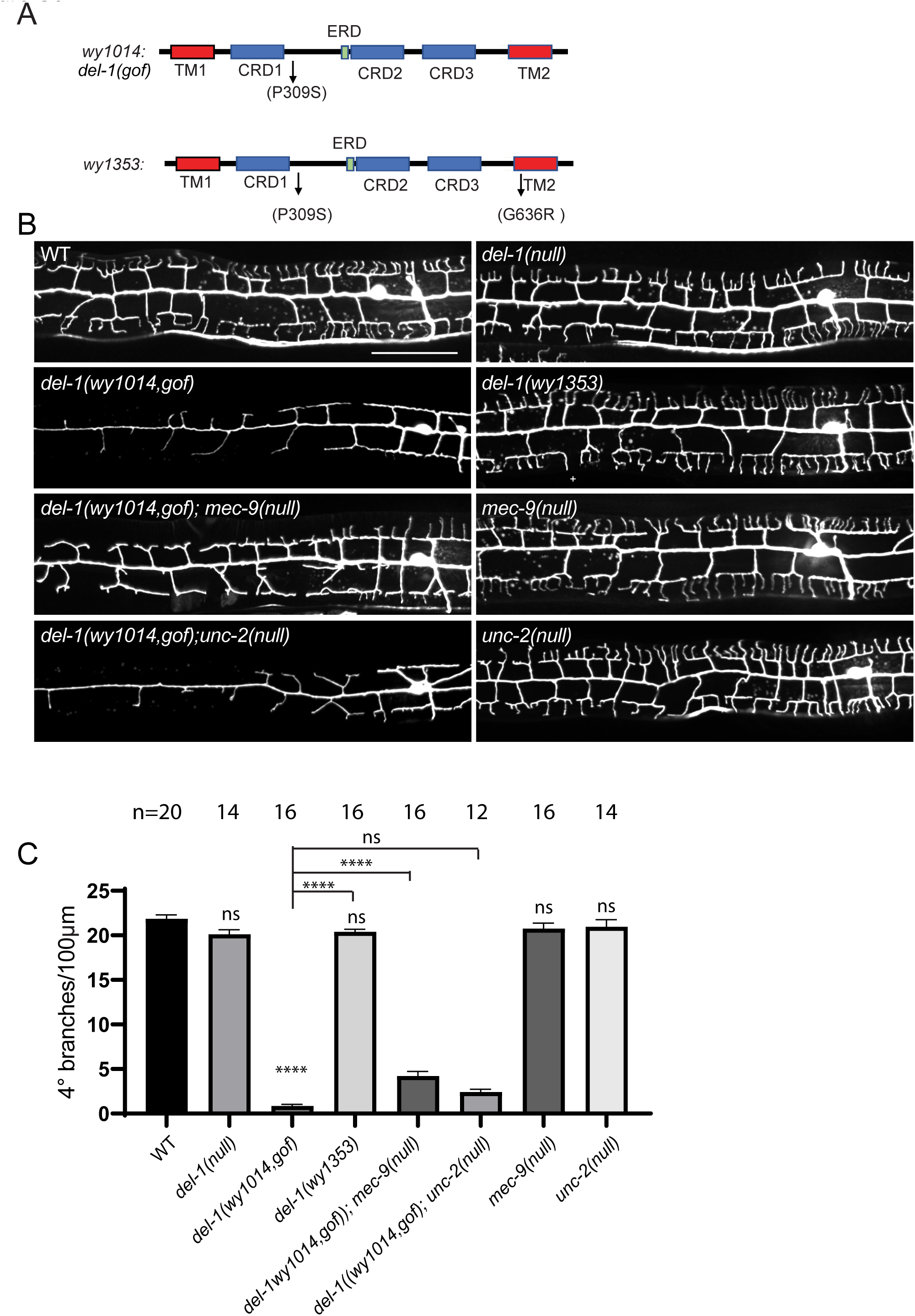
*wy1014* is a gain of function allele of *del-1,* related to Figure 5. (A) Illustration of DEL-1 isoform a with alleles indicated. CRD: Cysteine-rich domain. ERD: Extracellular regulatory domain. TM: transmembrane domain. (B) Representative confocal images of PVD morphology in the indicated genotypes at the L4 stage. Scale bar: 50 μm. (C) Quantification of the number of 4° dendrites per 100 μm the indicated genotypes. Statistics, ANOVA with a Tukey correction. ****p<0.0001. ns: not significant. Error bars: SEM.

**Fig. S7.**
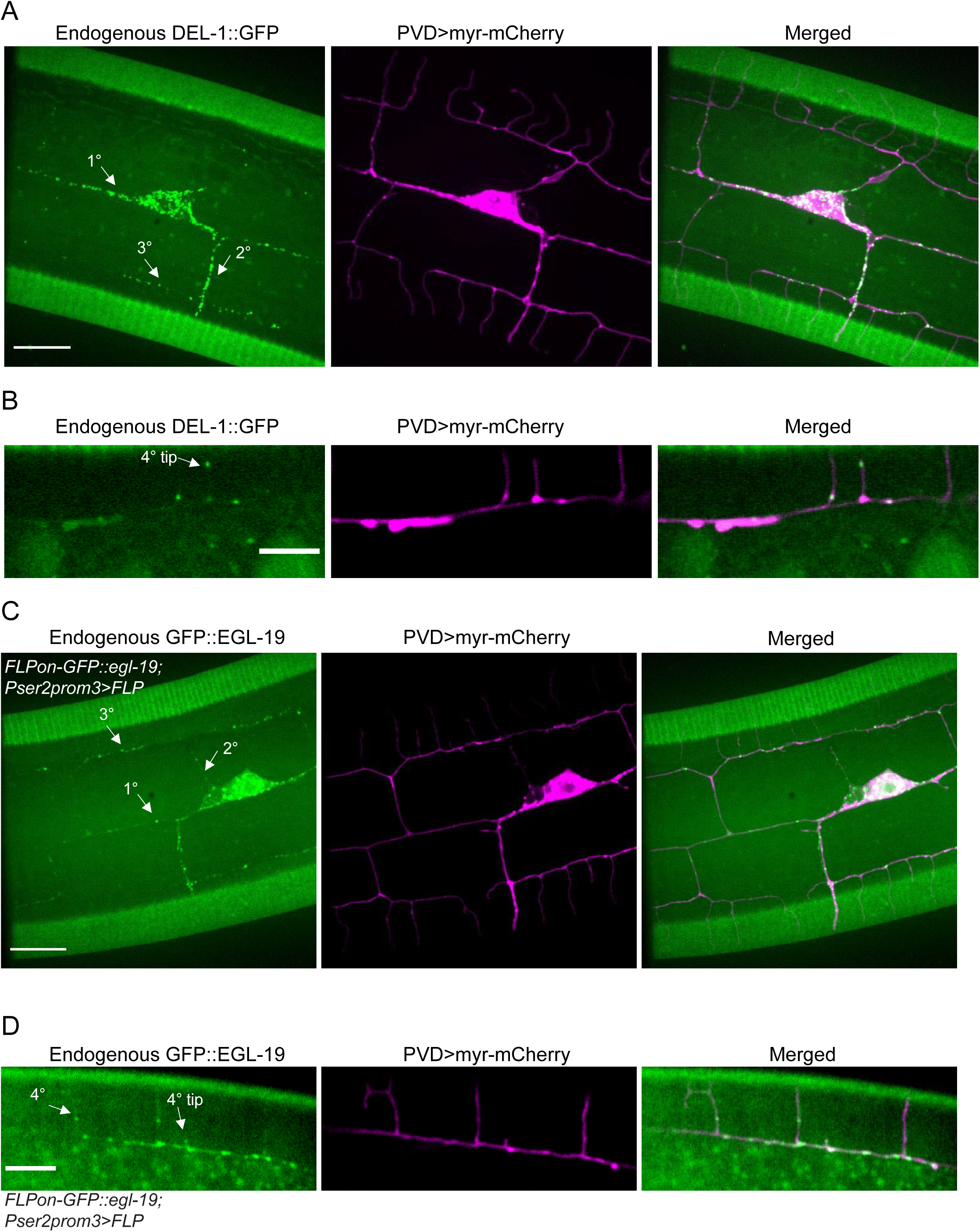
DEL-1 and EGL-19 are expressed in PVD and localize to the tip of 4° dendrites, related to Figure 5. (A and C) Confocal images showing the expression pattern of endogenously tagged DEL-1::GFP (A) and PVD cell-specific endogenously tagged GFP::EGL-19 (C) are expressed in PVD cells and localized along 1°, 2°, 3° dendrites. Scale bar: 10 μm. (B and D) Confocal images from a single z-plane showing endogenously tagged DEL-1 (B) and PVD cell-specific endogenously tagged EGL-19 (D) localized along or at the tip of 4° dendrites. Scale bar: 5 μm. All images are of early L4 stage worms.

**Fig. S8.**
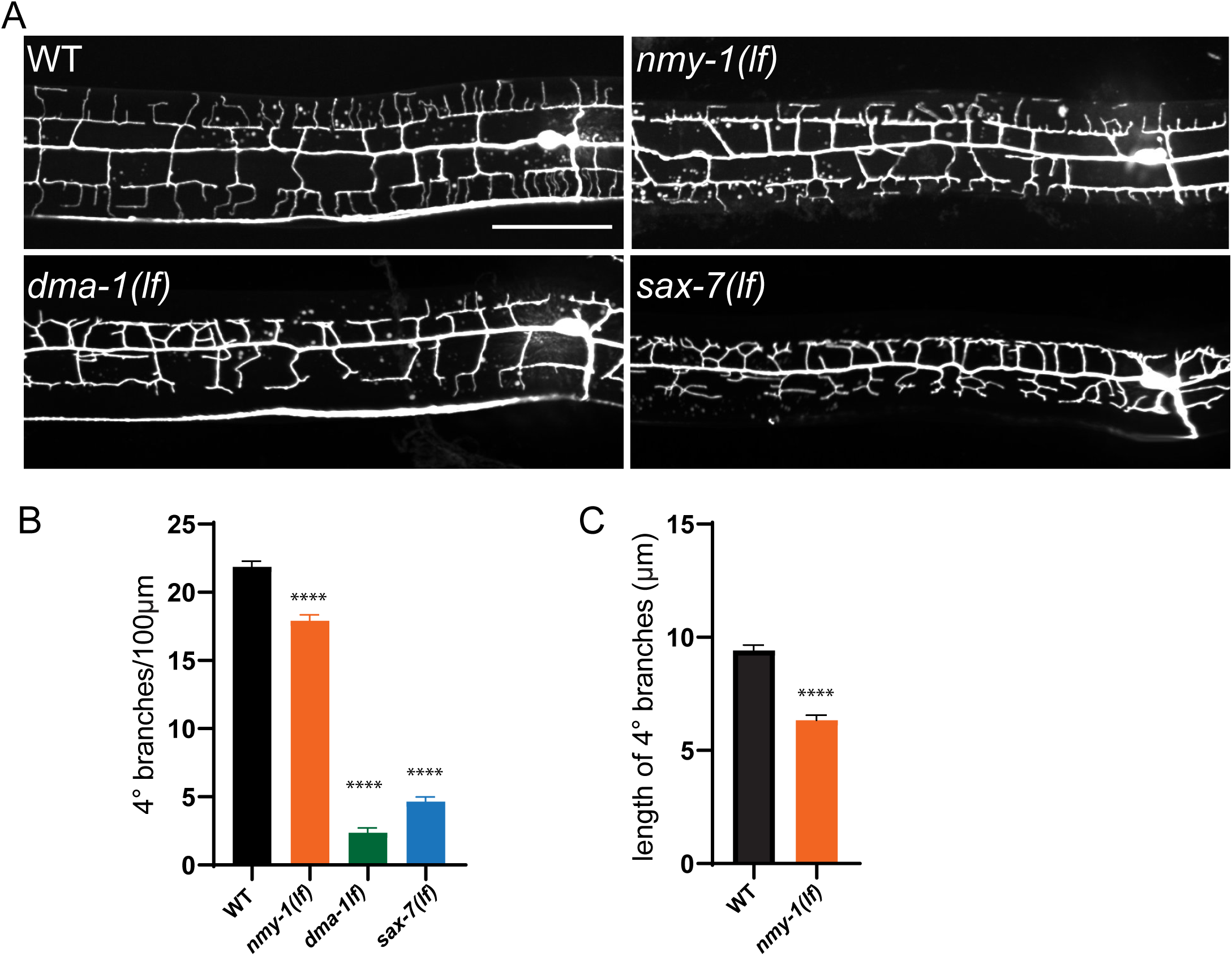
Weak loss of function alleles of *sax-7*, *dma-1* and *nmy-1* show fewer 4° dendrites, related to Figure 6. (A) Confocal images showing PVD morphology of indicated genotypes at the L4 stage. Scale bar: 50μm. (B) Quantification of the number of 4° dendrites per 100 μm in the indicated genotypes. n=20 for WT and *nmy-1(sb113)* mutant. n=13 for *dma-1(wy50544)* and *sax- 7(wy1127)* mutants. One-way ANOVA with a Tukey correction was used for statistical analysis. ****p<0.0001. Error bars: SEM. (C) Quantification of the average length of 4° dendrites in wild- type and *nmy-1(sb113)* mutants. 10 worms are quantified for each genotype. Student’s T-test was used for statistical analysis. ****p<0.0001. Error bars: SEM.

**Table S1.**
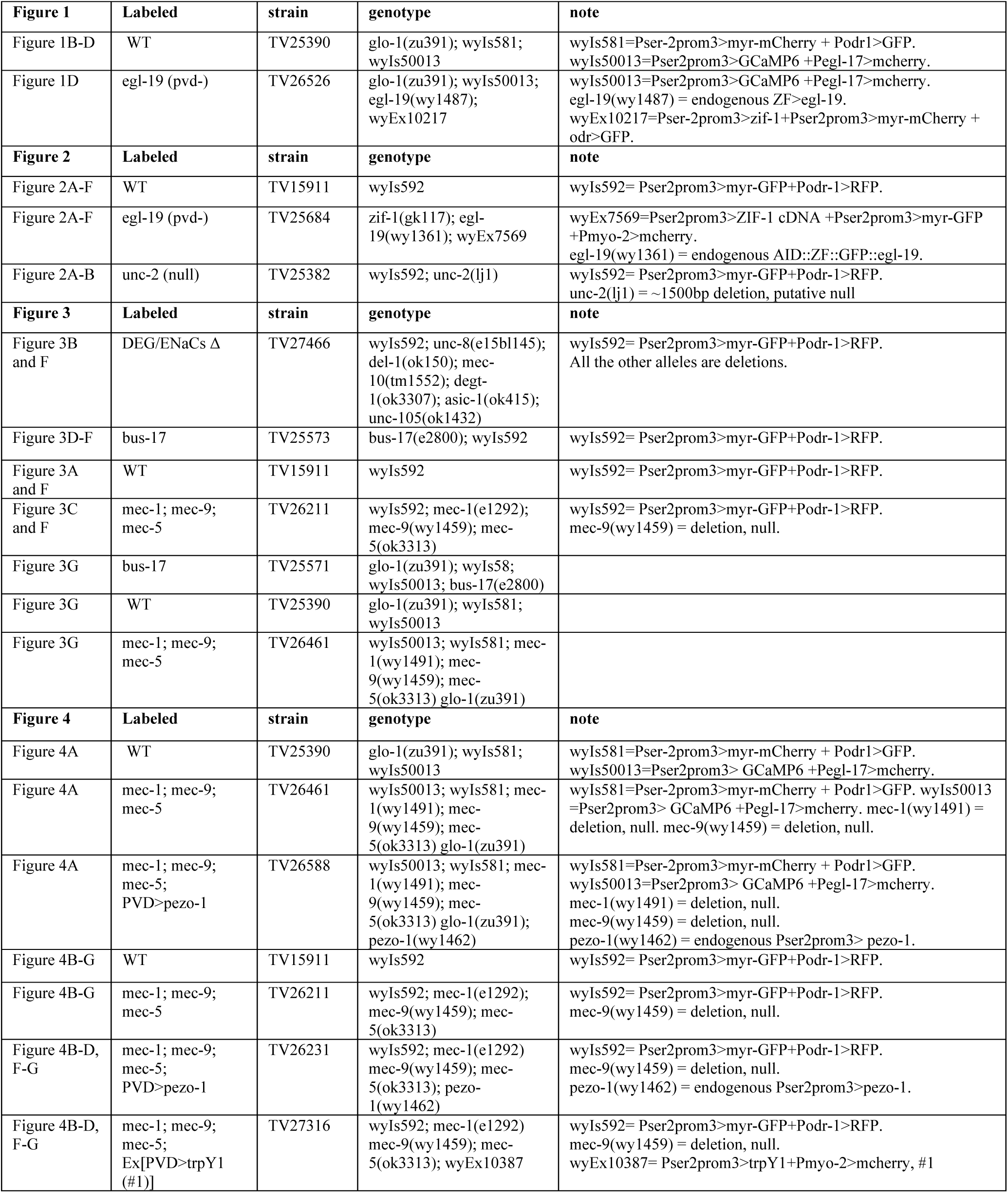
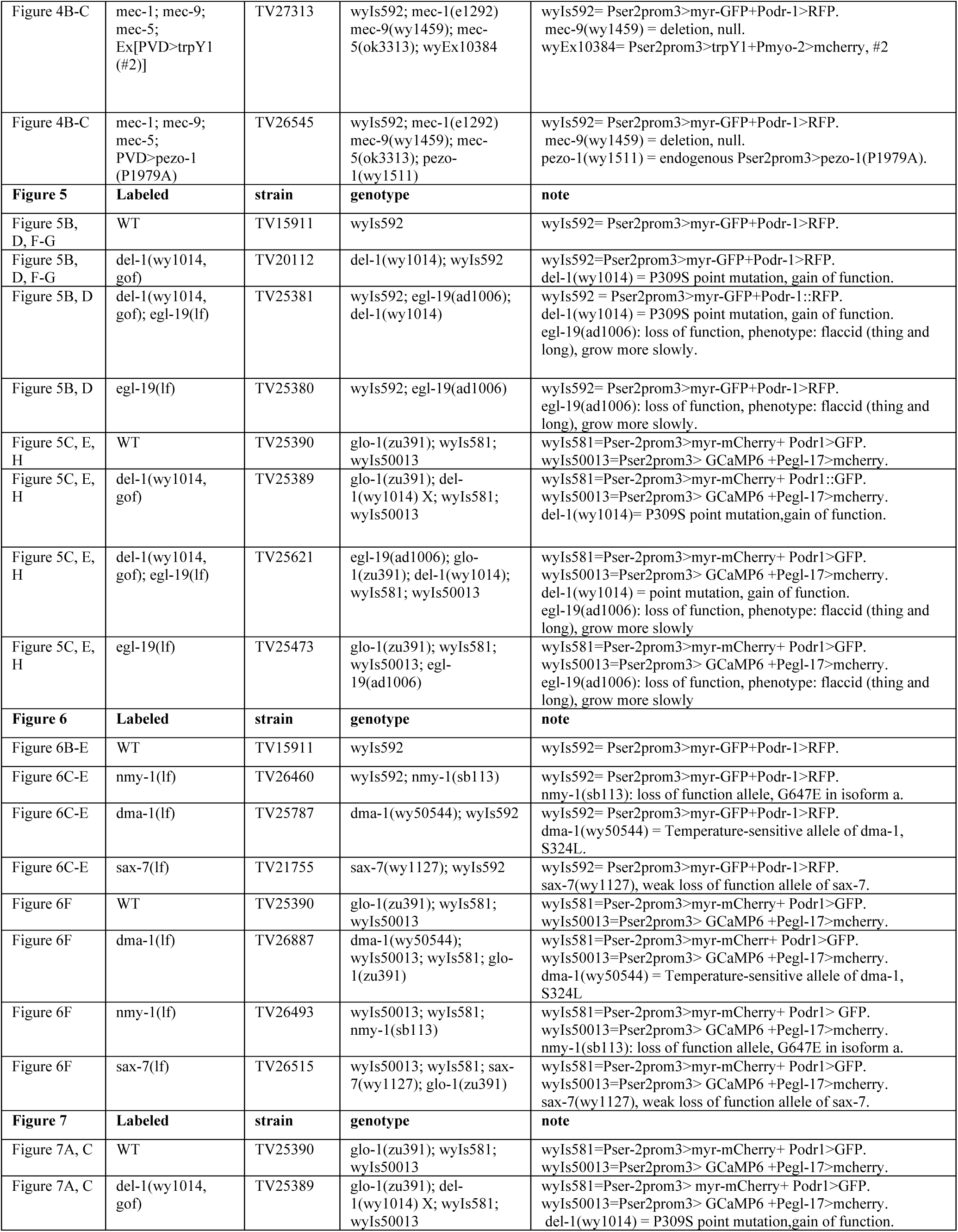
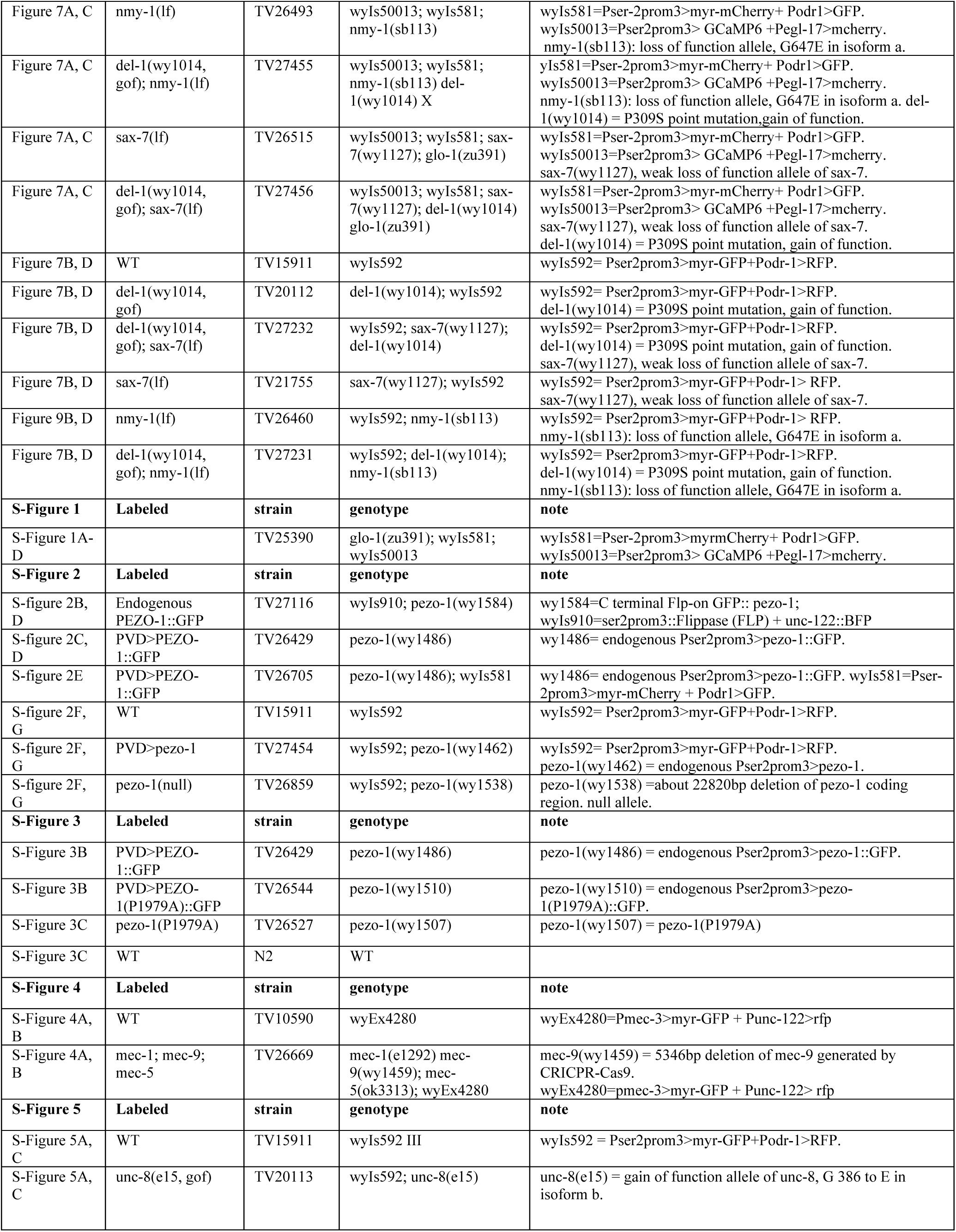
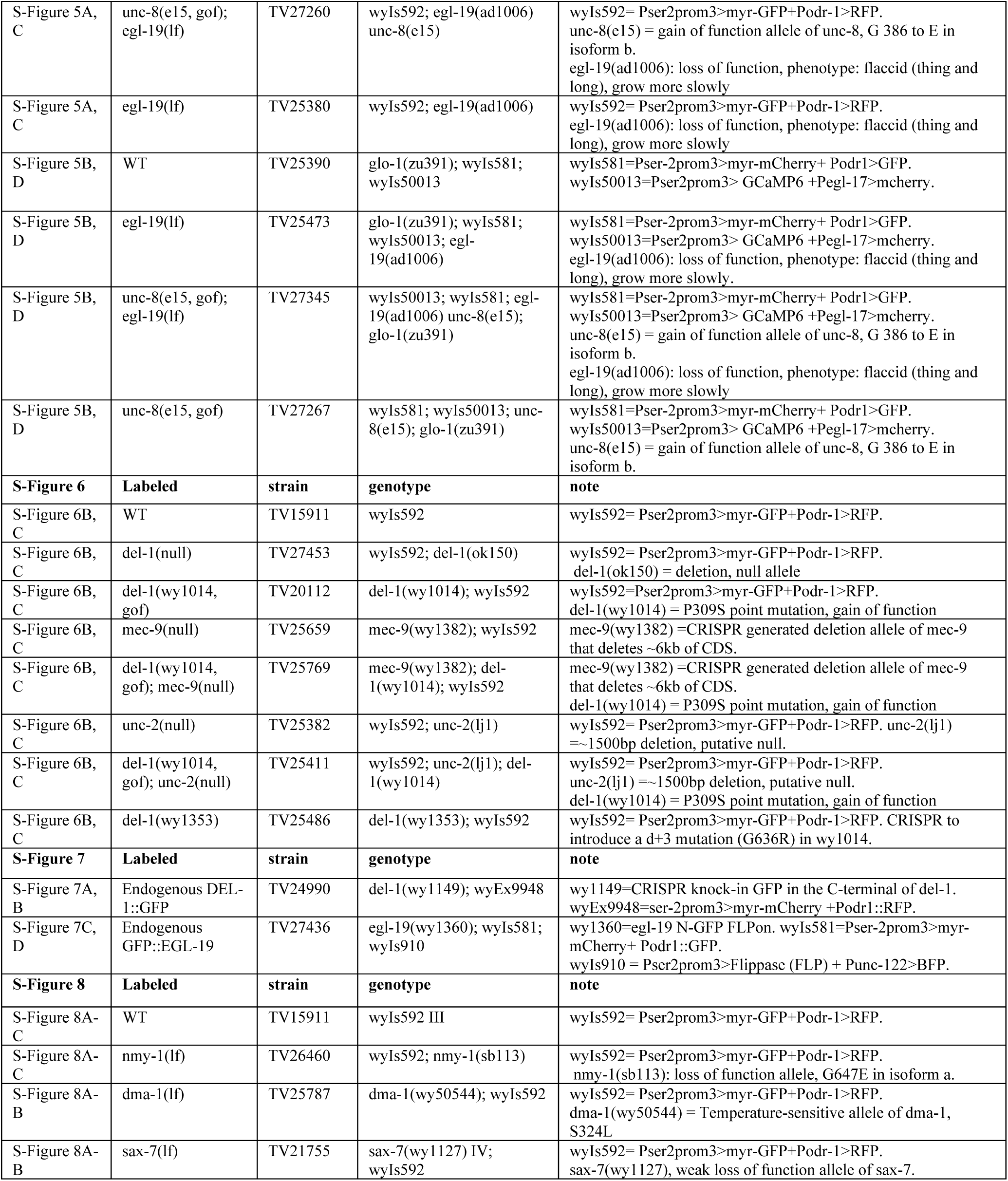
*C. elegans* strains.

**Table S2.**
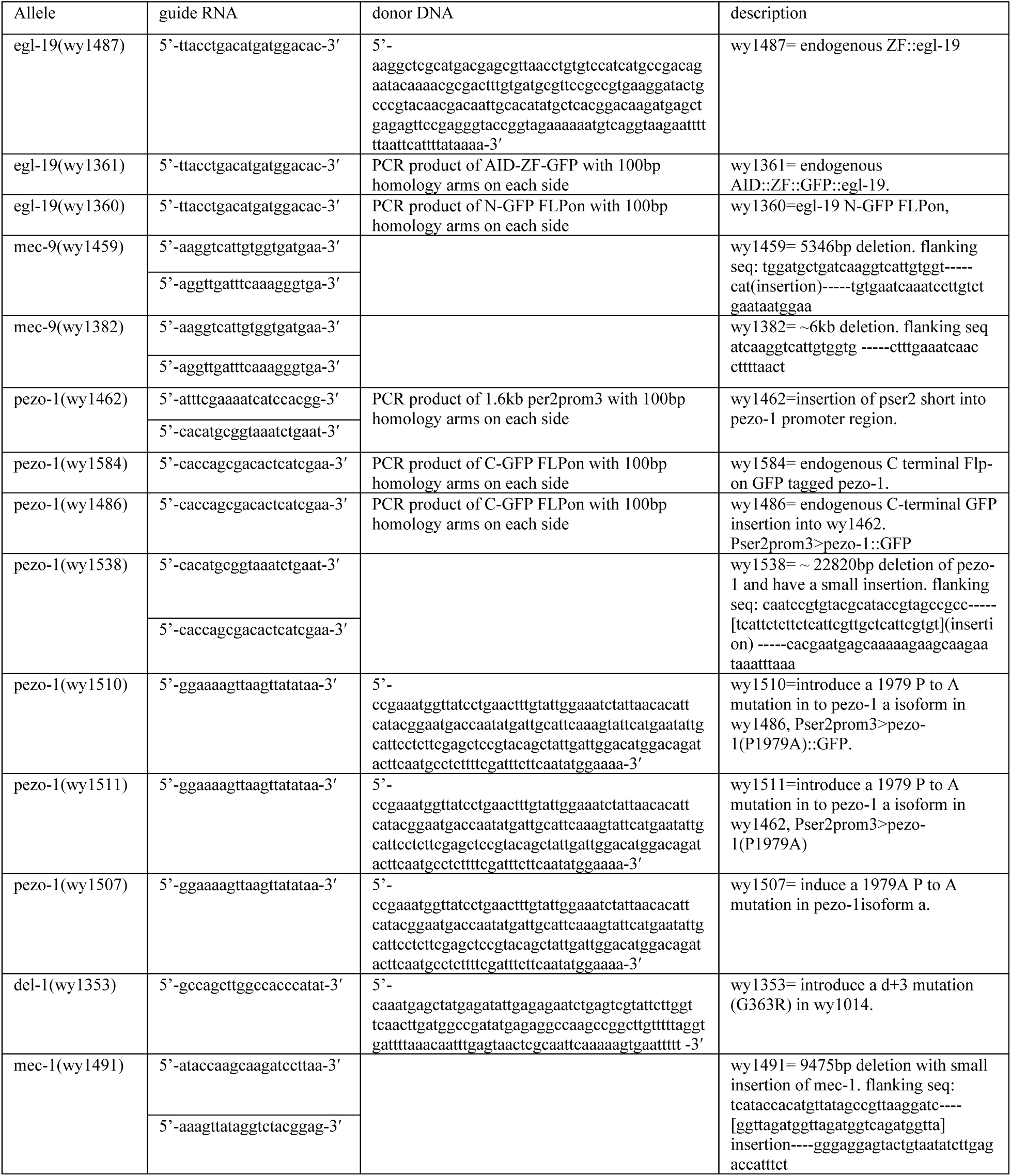
Alleles generated by CRISPR/Cas9-Mediated Genome Editing.

